# A lipid-associated macrophage lineage rewires the spatial landscape of adipose tissue in early obesity

**DOI:** 10.1101/2022.02.26.482134

**Authors:** Cooper M. Stansbury, Gabrielle A. Dotson, Harrison Pugh, Alnawaz Rehemtulla, Indika Rajapakse, Lindsey A. Muir

## Abstract

**Objective:** Obesity-induced metabolic dysfunction, tissue remodelling, and chronic inflammation in visceral white adipose-tissue (WAT) are correlated with insulin resistance, type II diabetes, and metabolic disease pathogenesis [1]. In this work, we sought to establish spatio-temporal context of adipose tissue macrophage (ATM) reprogramming during obesity.

**Methods:** We captured single-cell RNA-sequencing, spatial transcriptomics, and histological imagining of murine WAT over the course of diet-induced obesity to study macrophage phenotype dynamics. We developed a straightforward mathematical approach to integrating multi-modal data to quantify obesity-induced changes to WAT organization. We aligned ATM phenotypes with crown-like structures (CLS) in early obesity and used spatial network analysis to uncover signalling mechanisms implicated in CLS formation.

**Results:** We identified novel diversity of the lipid-associated macrophage (LAM) phenotype, whose transcriptional profile, signaling mechanisms, and spatial context serve as indicators of CLS formation in early obesity. We demonstrated that dysregulation of lipid-metabolic signalling is a critical turning point in the monocyte-LAM lineage and identified novel ligand-receptor mechanisms including *Apoe, Lrp1, Lpl* and *App* that serve as hallmarks of nascent CLS in WAT.

**Conclusions:** Multi-modal spatio-temporal profiling demonstrates that LAMs disproportionately accumulate in CLS and are preceded by a transition-state macrophage phenotype with monocytic origins. We identified novel ligand-receptor interactions implicated in nascent CLS regions which may guide future cellular-reprogramming interventions for obesity-related sequelae.

**Graphical Abstract:** 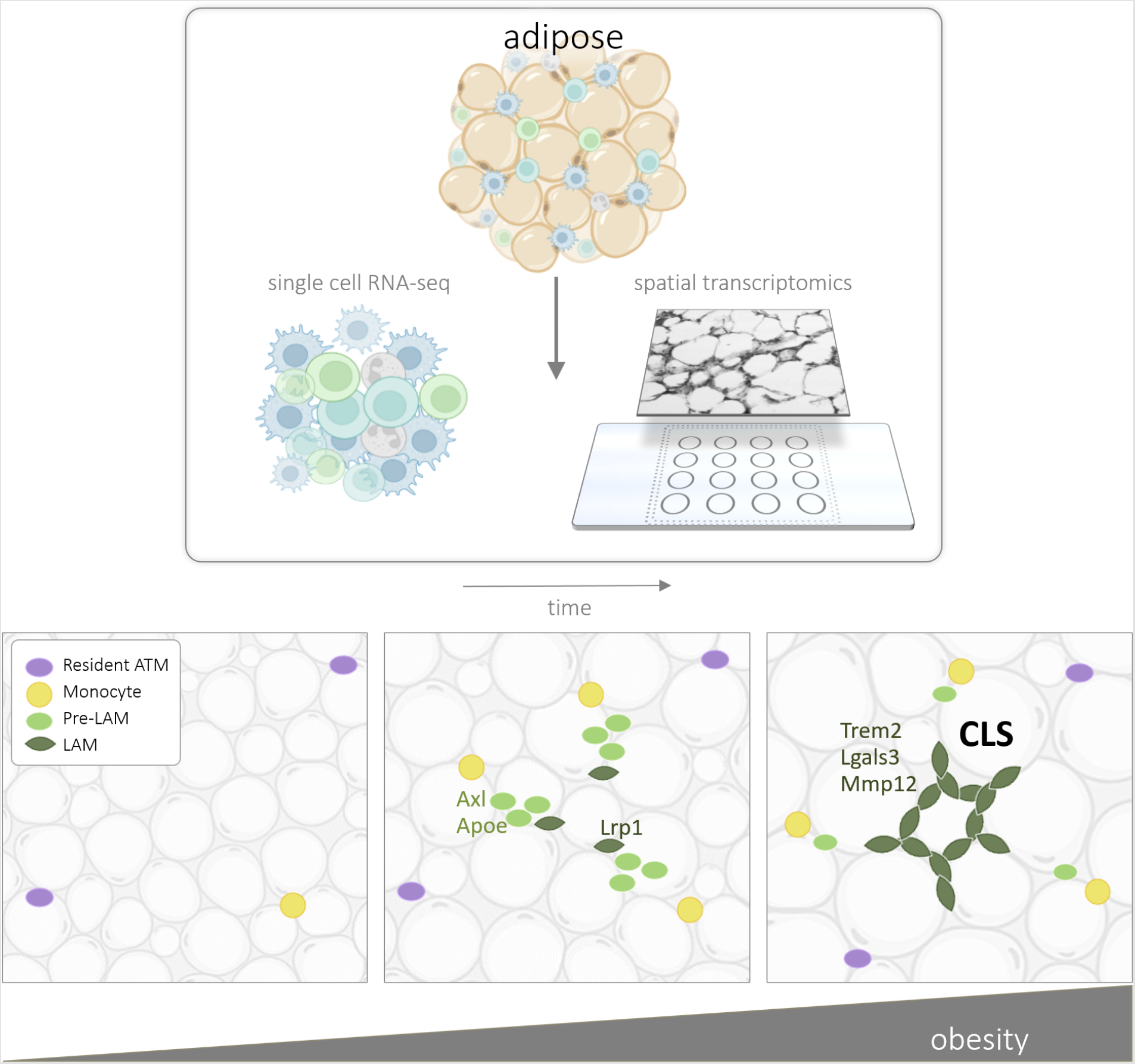

**Highlights:**

- We characterize a novel lipid-associated macrophage (LAM) phenotype along the monocyte-LAM lineage
- Integrated imaging, single-cell sequencing and spatial transcriptomics data show that LAMs accumulate at nascent CLS
- Analysis of spatial transcriptomics data reveals a novel set of ligands and receptors that implicate immature LAMs in shaping the CLS microenvironment in early obesity
- We present a simple mathematical framework for studying dynamics of tissue-structure over time

## 1. INTRODUCTION

Obesity is associated with chronic inflammation and metabolic dysfunction in mice and humans [2, 3, 4, 5]. Increased metabolic demand requires remodeling of white adipose-tissue (WAT) that results in changes to WAT structure and function [6, 7]. Normal WAT function requires coordination between multiple cell types including stromal vascular cells, immune cells, and adipocytes, which are the largest cellular constituent of WAT by volume [6, 8]. In obesity, WAT composition is dramatically altered and cells undergo dynamic changes to their morphology and phenotype that culminate in adipocyte hypertrophy and cell death [6, 9]. The dynamics of WAT immune cells during obesity are well-documented, but the molecular mechanisms regulating immune and metabolic dysfunction and their spatial organization within WAT remain poorly understood.

Immune cells help maintain healthy WAT homeostatic function and participate in WAT remodeling in response to changes in metabolic demand. The hallmark of obesity-induced immune dysregulation is increased abundance and diversity of macrophages in WAT [10, 11, 12]. Both tissue-resident macrophages and macrophages derived from recruited monocytes acquire poorly understood activation states during obesity-induced WAT remodelling [10, 11, 13, 14]. Changes in the macrophage transcriptional program are critical milestones in the development of insulin resistance, type II diabetes, and other metabolic disorders [10, 14] and are shown to persist after weight loss [15, 16, 12].

Previous single cell studies have cataloged WAT cellular composition, thus refining our understanding of immune cell phenotypes in obesity [10, 8, 13, 11]. However, single cell molecular profiling does not allow for analysis of the spatial patterning of tissue structure. Recent studies in humans have mapped single cell genomic profiles onto spatial transcriptomics data in order to characterize spatial patterning WAT cellular composition [6, 17]. However, a spatial understanding of obesity-induced WAT-remodelling over the timecourse of metabolic disruption is lacking.

We sought to spatially contextualize immune cell phenotype dynamics in early and chronic obesity. In this study we sequenced thousands of single cells from murine WAT at different stages of diet-induced obesity and characterized transcriptional dynamics associated with the development of insulin resistance. To characterize the spatial context of obesity-driven immune cell dysregulation, we mapped tissue-specific genomic signatures to the WAT landscape using spatial transcriptomics. We developed a network approach to analyze the spatial organization of immune-dysregulation and used graph-theoretic measures to quantify changes to WAT structure.

We quantified the spatio-temporal dynamics of WAT macrophage infiltration and differentiation and identified cellular signalling mechanisms implicated in WAT remodelling. We describe novel diversity of the *Trem2* ^+^ lipid-associated macrophage (LAM) phenotype, whose transcriptional profile, molecular signalling mechanisms, and spatial context suggest a critical role in the formation of CLS in early obesity.

## 2. MATERIALS AND METHODS

### 2.1. Animals

C57BL/6J mice were used for all experiments (Jackson Laboratories 000664). Male mice were fed ad libitum a control normal chow diet (ND; 13.4% fat, 5L0D LabDiet) or high-fat diet (HFD; 60% calories from fat, Research Diets D12492) for the indicated amount of time starting at 9 weeks old. Animals were housed in a specific pathogen-free facility with a 12 h light/12 h dark cycle and given free access to food and water except for withdrawal of food for temporary fasting associated with glucose tolerance tests. All mouse procedures were approved by the Institutional Animal Care and Use Committee (IACUC) at the University of Michigan (Animal Welfare Assurance Number D16-00072 (A3114-01), #PRO00008583), and care was taken to minimize suffering adhering to the Institute of Laboratory Animal Research Guide for the Care and Use of Laboratory Animals.

### 2.2. Glucose Tolerance Tests

For glucose tolerance tests (GTT), starting four hours into the light cycle, mice were fasted with ad libitum access to water for six hours in clean cages. A 100 mg/mL D-glucose (Sigma G7021) solution was prepared in sterile -/- DPBS and injected at 0.7 g/kg of body weight. Area under the curve (AUC) calculations were performed using the log trapezoidal method.

### 2.3. Stromal Cell Isolation and Immune Cell Enrichment

Stromal vascular cells (SVCs) were collected from adipose tissues as in [4]. After cardiac perfusion, adipose tissues were collected, minced finely to 3-5 mm pieces, and added to ice cold HBSS+Ca/Mg. Up to 1.5 g of tissue per sample was digested in 10 ml of 1 mg/mL collagenase II (Sigma C68850) in HBSS+Ca/Mg at 37^°^ C for 45 minutes with vigorous shaking. Digests were filtered through buffer-soaked 100 micron cell strainers and centrifuged at 300 x g at 4C to pellet SVCs. SVCs were enriched for CD45^+^ immune cells using Biolegend MojoSort Mouse CD45 Nanobeads (Biolegend 480027), following the manufacturer’s protocol. Briefly, SVC pellets were resuspended in 1 mL MojoSort Buffer, pooling the four samples from each cohort into a single respective cohort tube (ND, 8w, 14w), then filtered through a 70 micron cell strainer and placed in 5 mL polypropylene tubes. After addition of nanobeads, samples were sequentially processed for magnetic separation. Three magnetic separations in total were performed on the labeled fractions for increased purity. Final cell suspensions were filtered through 40 micron pipette tip filters. Cell viability was >80% with <15% aggregation.

### 2.4. Feature Barcoding and Single Cell RNA-sequencing Library Preparation

CD45^+^ SVCs were feature barcoded using TotalSeqB (Biolegend) antibodies (F4/80, CD11b, Mac-2, CD3, CD4, CD19). Library preparation was performed by the University of Michigan Single Cell Sequencing core using the 10x Genomics Chromium Single Cell Kit (3’V3, #220103/PN120262). 100 million reads from up to 5,000 cells were collected for single cell transcript data, and 25 million reads from up to 5,000 cells were collected for feature barcoding data.

### 2.5. Spatial transcriptomics tissue and library preparation

Within 30 minutes of cardiac perfusion, epididymal WAT samples that were contralateral to those used for scRNA-seq were pre-soaked in ice cold O.C.T. Compound (VWR 25608-930) and placed in biopsy cryomolds (VWR 25608-922) with fresh O.C.T., rapidly frozen by immersion in isopentane cooled using liquid nitrogen, and kept on dry ice or at -80°C until sectioning. Fresh tissue sections were cut at 10 *μ*m after 20 minute equilibration in a cryochamber set to -26^°^C or below with specimen arm at -40^°^C. Sections were placed onto the Visium Spatial Gene Expression slide and subsequent processing and library preparation were performed by the University of Michigan In Vivo Animal Core pathology laboratory and the Advanced Genomics Core according to the manufacturer’s protocol (10x Genomics PN-1000184).

### 2.6. Tissue histology and immunostaining

Hematoxylin and eosin (H&E) and immunostaining were performed in the ULAM In Vivo Animal Core pathology laboratory at the University of Michigan. After fixation for 48 hours in 10% neutral buffered formalin, tissues were trimmed, cassetted, and processed to paraffin in an automated tissue processor (TIs-sueTek, Sakura). Processed tissues were embedded in paraffin and sectioned at 4 microns on a rotary microtome (Leica Biosystems, Buffalo Grove, IL). Tissues were mounted on glass slides and stained with hematoxylin and eosin using routine protocols on an automated histostainer (Leica ST5010 Autostainer, Leica Biosystems), followed by coverslipping.

### 2.7. Data processing

Single cell RNA-sequencing files were processed using the 10X Genomics CellRanger (version 4.0.0) pipeline. The resulting filtered matrices were analyzed using `scanpy` [18]. Briefly, we filtered out cells that did not express at least 500 genes and genes that were not expressed in at least 10 cells, resulting in 13,820 cells and 31,053 genes across all diet conditions (1,261 ND cells, 6,123 8w HFD cells, and 6,436 14w HFD cells). We normalized read-counts per cell after filtering. Spatial sequencing data were processed using the 10X Genomics SpaceRanger (version 1.0.0) pipeline with mouse reference GRCm38, and resulting feature-barcode matrices were loaded into `scanpy` [18] for further analysis. We filtered out capture spots that expressed fewer than 5 genes from all subsequent analysis. We normalized read-counts per capture spot after filtering.

### 2.8. scRNA-seq clustering and visualization

Clustering was performed on cells from each time point independently using Algorithm 1. Preprocessing and clustering were performed using Python and the single cell gene expression package `scanpy` [18]. scRNA-seq data were normalized and log-transformed before dimension reduction using principal component analysis (PCA) with *r* = 50. We constructed the similarity matrix **A** using *k* = 9 neighbors and Euclidean distance prior to clustering with the Leiden clustering method [19] with resolution parameter *γ* = 0.95. This analysis resulted in 18 clusters in ND, 25 in 8w HFD fed mice, and 20 in 14w HFD fed mice. Visualization of data was performed using uniform manifold approximation and projection (UMAP) [20]. Dimensionality was reduced using PCA (*r* = 10) on the combined set of genes with non-zero expression at all three time-points. Cells were passed to UMAP with the following parameters: `n_neighbors=50`, `min_dist=0.25` and `metric=‘euclidean’`.

#### Algorithm 1

Clustering and Visualization

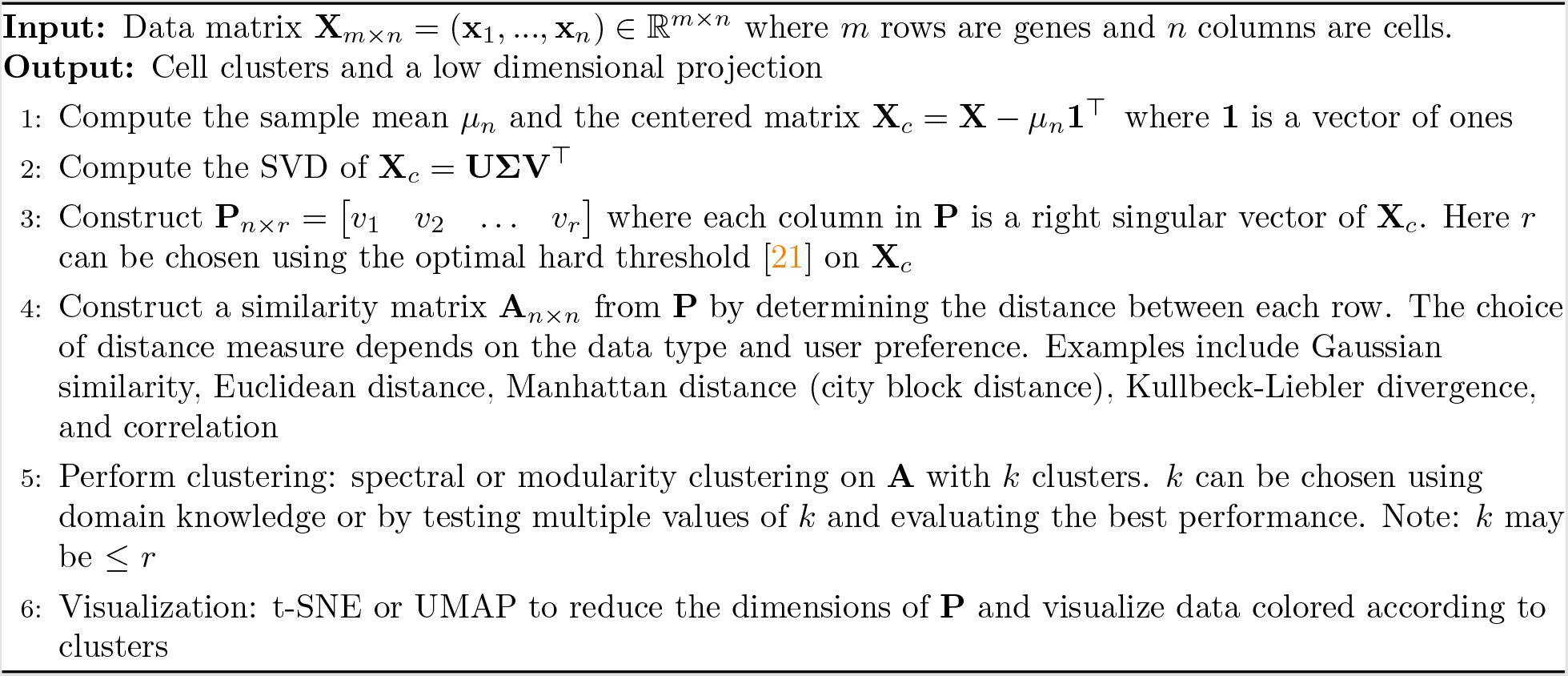

### 2.9. scRNA-seq cell type annotation

Annotation of cell types after clustering was performed using ranked expression of cell-type specific mouse markers genes from PanglaoDB [22]. The top 50 most unique marker genes were used for each cell type sorted by their ubiquitousness index. Each cluster was assigned to a cell type based on the maximum mean rank of marker genes amongst the differentially expressed genes for that cluster. A small set of 165 CD45^+^ cells were also identified that did not align with major immune cell populations; this population was excluded from subsequent analyses. We performed differential expression analysis on clusters and sorted genes by their Student’s T-Test statistic computed using the `scapny.tl.rank_genes_groups()` function with `method=‘t-test’`.

### 2.10. Mapping cell-type signatures to spatial transcriptomics data

We used a conditional autoregressive-based deconvolution (CARD) model (https://github.com/YingMa0107/ CARD) to spatially deconvolute cell type signatures of our data and estimate the strength of cell type proportions across tissue capture spots [23]. CARD was chosen over other deconvolution methods for its ability to leverage nearby spatial information during cell type proportion estimation using a conditional autoregressive modeling assumption, which imposes spatial correlation structure on the outputs. Briefly, each single cell was annotated for cell type and scRNA-seq count matrices and spatial transcriptomics count matrices were structured according to CARD documentation. Deconvolution was performed using `createCARDObject()` with parameters `minCountGene=10`, and `minCountSpot=20`. Outputs were stored as tabular files for downstream analysis. CARD estimates the cell type proportions for *k* cell types defined given *g* genes at *n* tissue spots using the following non-negative matrix factorization model:

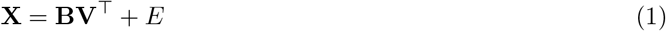

where **X** ∈ ℝ ^*g*× *n*^ is the spatial transcriptomics data matrix, **B** ∈ ℝ ^*g* × *k*^ is a matrix of aggregate cell type signatures derived from the scRNA-seq data, **V** ∈ ℝ ^*n* × *k*^ is a matrix of cell type proportions at each tissue spot and E ∈ ℝ ^*g* × *n*^ is a normally distributed error matrix. For further details, see [23].

### 2.11. Macrophage continuum analysis

A linear model was used to quantify cells along a user-defined continuum as in [24] and [25]. The procedure from [24] is generalized in Algorithm 2. Briefly, we used Ordinary Least Squares (OLS) to linearize the correlation between two states of interest in a given cell population, e.g., ATM-LAM or monocyte-LAM. We quantified each cell’s position relative to the states of interest by computing the distance between the cell and the each state along the OLS solution. We defined a gene set using DE between the two states with a Bonferroni correction for multiple-tests to *α* = 0.05 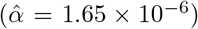 and chose top genes for each pole, ranked by their fold change.

#### Algorithm 2

Continuum Quantification

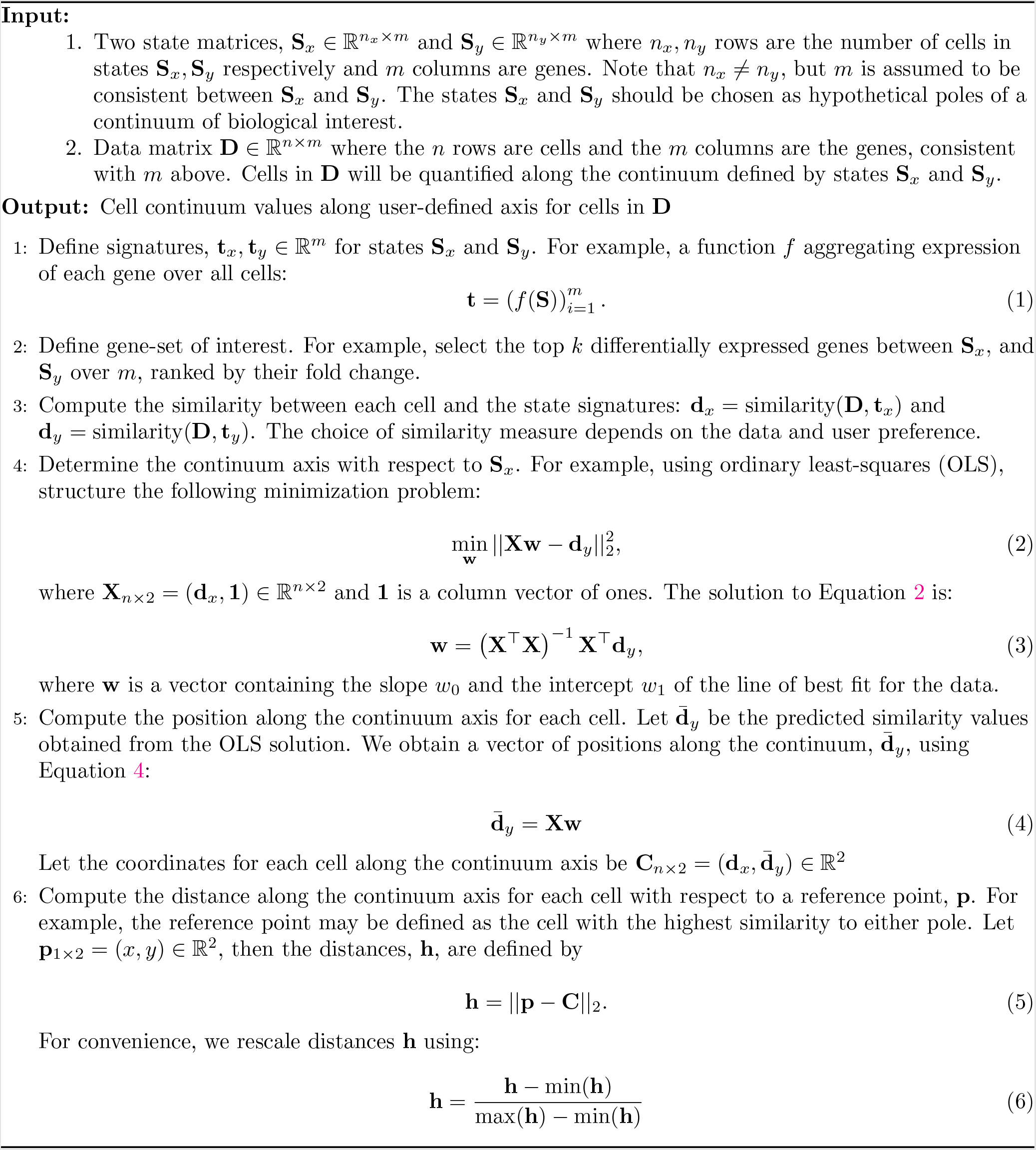

### 2.12. Ligand-receptor colocalization

We obtained mouse ligand-receptor (LR) pairs from [1]. We defined colocalization the simultaneous expression of ligand, l and receptor r at a given tissue-capture spot t. The colocalization ‘strength’ or l and r at t was quantified using the geometric mean of normalized expression:

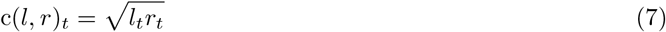

Where *l*_*t*_ and *r*_*t*_ are the expression of *l* at *t* and *r* at *t* respectively. By using the geometric mean we ensure that *c(l, r)* = 0 where either *l*_*t*_ = 0 or *r*_*t*_ = 0. LR pairs are said to be colocalized wherever *c*(*l, r*)_*t*_ > 0. Time-dependent colocalization between LR pairs was taken as a necessary, but not sufficient condition in determining possible signalling pathways. We computed the proportion of spots where *l* and *r* were localized and normalized the proportion to 1k spots to account for differences in tissue-section sizes.

### 2.13. Construction and analysis of network models

We aim to construct a network model that preserves spatial relationships in tissue structure. Let **G** be a finite, simple, and undirected graph with node set *V* (**G**) = {1, 2,. .., *n*} and edge set *E*(**G**) ⊂ *V* (**G**) × *V* (**G**). Let *e*_*ij*_ be an edge between node *i* and node *j*. The n nodes of **G** are chosen from the set of tissue-capture spots from the spatial transcriptomics data matrix. Thus, each node *i* has a specified spatial position in a 2-dimensional Euclidean plane, *p*_*i*_ ℝ^2^. Edges are defined between nodes as a function of (1) their Euclidean distance and (2) their nodal properties determined by the biological question of interest. In the simplest case, we may define a radius, *r*, which is the maximum physical interaction distance between two nodes. The strength of the relationship between node *i* and node *j* is encoded in the edge weight *w*_*ij*_. Edge weights are defined by a function, *f* : *V* (**G**) × V (**G**) → ℝ.

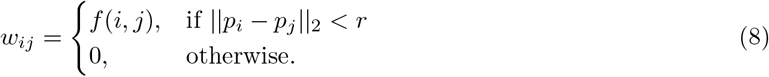

A network defined this way captures the spatial patterning of *f* in the local neighborhood constrained by *r*. It is also useful to define the weighted adjacency matrix of **G** to be the *n* × *n* matrix **A**(**G**) with rows and columns indexed by *V* (**G**). We will denote **A**(**G**) as **A** and the entry (*i, j*) of **A** as **A**(*i, j*) = *a*_*ij*_. The weighted adjacency matrix may be defined:

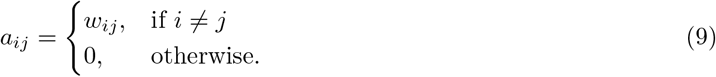

For example, we define LAM-networks based on the harmonic mean of Mac5 CARD estimated proportions over neighboring tissue spots [23]. In this case, the choice of the harmonic mean is based on the interpretation of CARD outputs as proportions of the tissue spot explained by a given cell type signature [23]. Let *m*_*i*_ be the proportion of Mac5 cell type at tissue spot *i*:

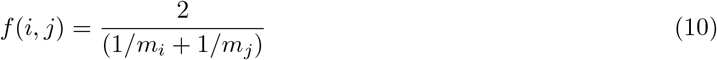

The concept of network centrality is motivated by identification of ‘important’ nodes of a network [26]. We focus on two measures of network centrality: degree centrality (Equation 11) and eigenvector centrality (Equation 12). Degree centrality is a ‘local’ measure of connectivity whereas eigenvector centrality is a ‘global’ measure of centrality. Let 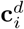 denote the degree centrality of node *i*. Degree centrality is the sum of all the edge weights of node *i*,

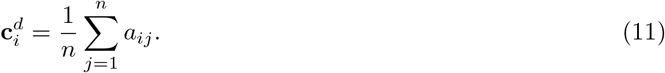

The eigenvector centrality of each node, defined here up to a scale factor, is proportional to the sum of the eigenvector centralities of its neighbors, that is:

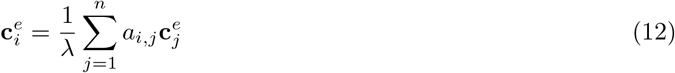

where **c**^*e*^ is an eigenvector of **A** and λ is the corresponding eigenvalue. The centrality is taken to be an eigenvector that corresponds to the largest eigenvalue of **A**.

### 2.14. Adipocyte sizing

Images of H&E stained adipose tissue (Materials and Methods: Tissue histology and immunostaining) were analyzed for adipocyte size using the Python package `skimage` [27]. Briefly, images were converted to greyscale and subjected to an unsharp masking filter with parameters: `amount=75` and `amount=100`. Filtered images were filtered again using a median filter with default parameterization followed by morphological reconstruction using `method=‘erosion’` to enhance contrast between neighboring cells. Finally, images were filtered using a Gaussian kernel with `simga=3`. Processed images were thresholded at the 25^th^ percentile before segmentation using the Watershed method. Properties of each segmented cell were obtained using `measure.regionprops()`. We computed the circularity, *C* of all segmentation using Equation 13.

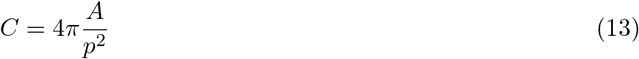

Where *A* is the estimated area and *p* is the estimated perimeter of the segmented cell. We filtered regions with 0.4 < *C* < 0.9 and regions with areas above or below 2.32 *σ* from the time-dependant mean.

### 2.15. Histological crown-like structure quantification

Tissue images captured during spatial transcriptomics tissue preparation (Materials and Methods: Spatial transcriptomics tissue and library preparation) were analyzed using a segmentation algorithm to classify each pixel into one of four categories: CLS_hi_, CLS_mid_, Other, and Adipocyte based on 3-channel pixel intensity values. Briefly, we used the Python package `skimage` to perform Multi-Otsu Thresholding on the 14 week RGB image tensor [27]. We then extracted basic features using `feature.multiscale basic features()` with the following parameters: `intensity=True`, `edges=False`, `texture=True`, `sigma min=1`, and `sigma max=16`. We developed a Random Forest segmentation model with 50 estimators using the Python package `sklearn`. We then used the segmentation model to analyze the remaining diet conditions. Regions surrounding spatial capture spots were segmented, and the proportion of pixels in each category were computed and compared.

## 3. RESULTS

### 3.1. Dynamic remodeling of adipose tissue is concurrent with glucose intolerance in early obesity

Our model of diet-induced obesity included mice fed a normal chow diet (ND) or a 60% high fat diet (HFD) for 8 or 14 weeks. HFD feeding increased body weight and epididymal white adipose tissue (eWAT) mass as expected (Figure 1B-D). Mean adipocyte area and frequency of large adipocytes increased at 8 and 14 weeks (Figure 1G-H, Methods 2.14). Glucose tolerance tests showed increased area under the curve (AUC) starting at week 1, with the largest AUC and variability at weeks 7 and 8 (Figure 1E-F), suggesting development of insulin resistance.

**Figure 1:**
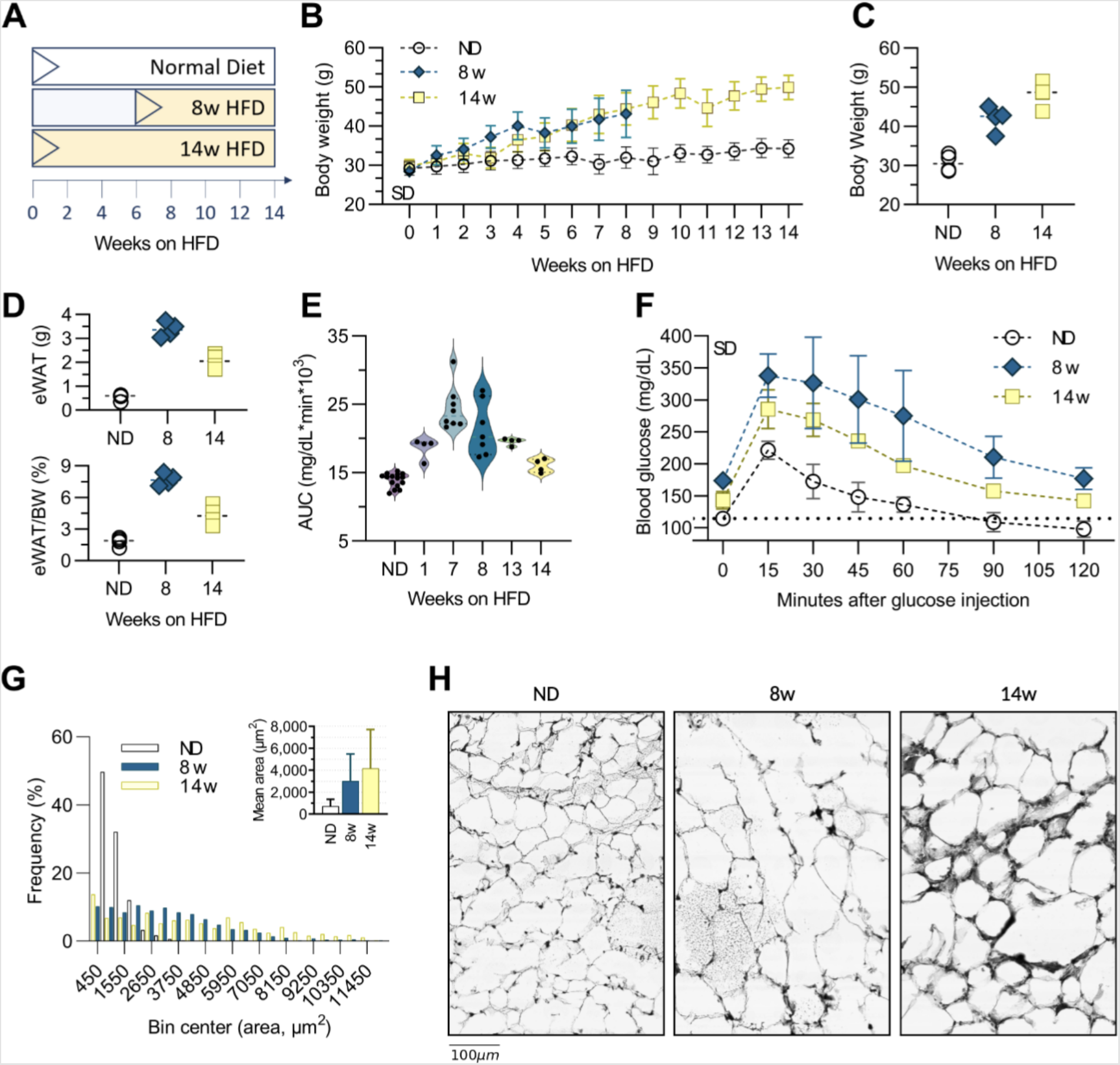
Diet-induced obesity and adipose tissue remodeling. **(A)** Time course for mice fed a 60% high-fat diet (HFD) for 8 weeks (8w) or 14 weeks (14w), versus normal diet (ND) controls. **(B)** Total body weight by week on HFD. **(C)** Final body weight at time of tissue collection. **(D)** Epididymal adipose tissue (eWAT) weight (top) and eWAT as a percentage of body weight (bottom). **(E)** Glucose tolerance test data showing area under the curve (AUC). **(F)** Glucose measurements for cohorts one week prior to endpoint tissue collection. **(G)** Frequency distribution and average adipocyte size in eWAT of ND, 8w, and 14w cohorts. **(H)** H&E images of adipose tissue sections at ND, 8 and 14 weeks on HFD.

### 3.2. Single cell profiling

It is well established that obesity induces changes in adipose tissue immune cells [10, 8], including accumulation of ATMs that promote metabolic dysfunction [2, 3]. However, the dynamics of these phenotypes remain incompletely understood. To examine immune cell dynamics in early and chronic obesity we performed single cell RNA-sequencing (scRNA-seq) on CD45^+^ cells from perigonadal (eWAT) fat pads of mice fed ND or fed a HFD for 8 or 14 weeks (n=4 per cohort).

Clustering and annotation of 13,820 single cells identified six broad immune cell populations: monocytes, T cells, B cells, dendritic cells, adipose tissue macrophages (ATM), and natural killer (NK) cells (Figure 2A), Methods Section 2.8). Antibody feature barcodes for select surface proteins that were used with scRNA-seq confirmed immune cell annotations (Figure S3, Methods Section 2.4). Annotations were additionally confirmed by comparison to cell type-specific gene expression profiles from public databases and published single cell genomic datasets (Figures S4-S6).

**Figure 2:**
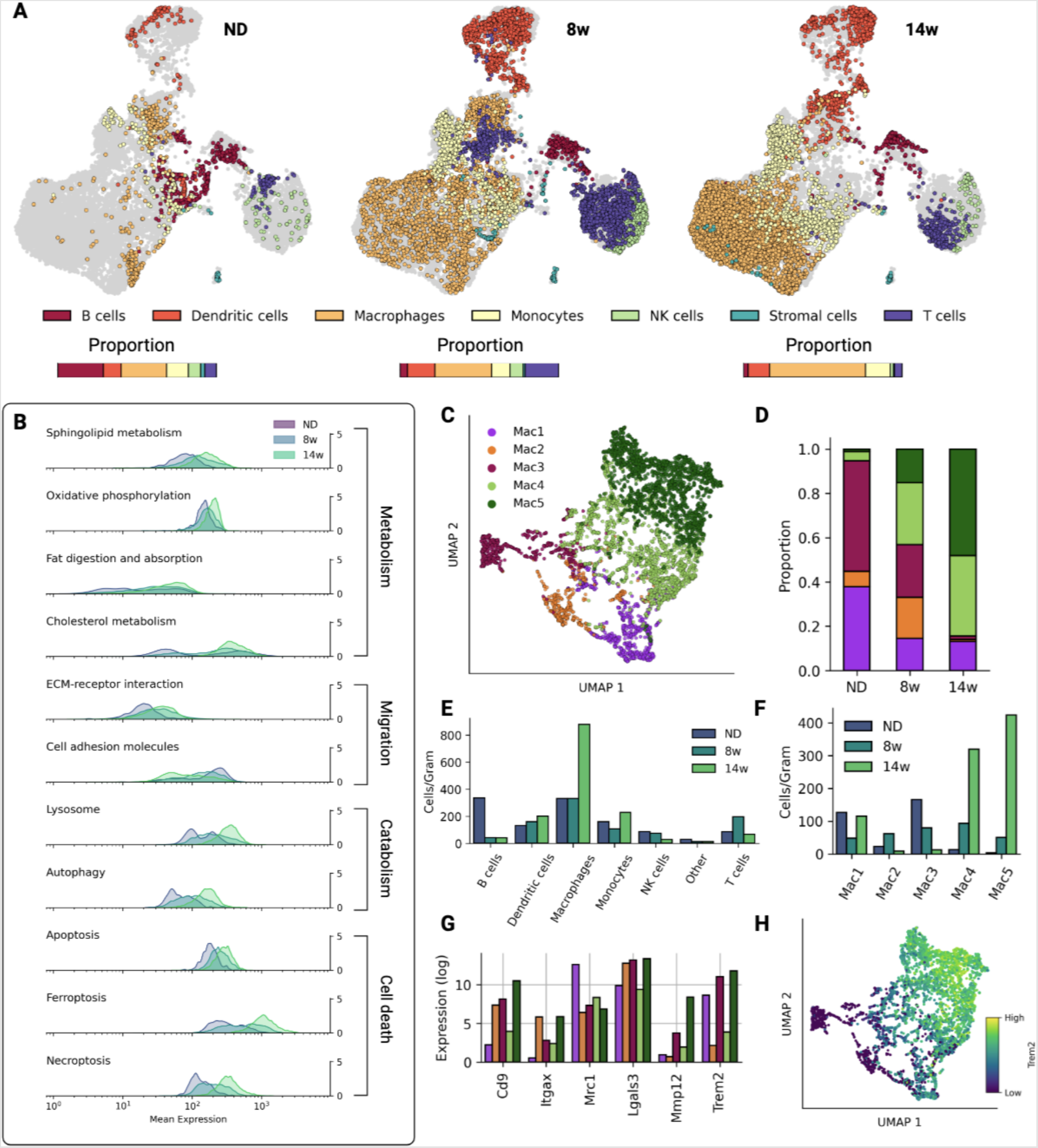
Single cell data on macrophage phenotypes in obesity. **(A)** immune cell population changes over the course of diet-induced obesity. **(B)** Changes in expression in expression of genes in select KEGG pathways in the macrophage subpopulation. **(C)** UMAP visualization of ATM clusters from scRNA-seq data. **(D)** Proportions of each ATM cluster at each time point. **(E)** The number of cells per gram of adipose tissue for each cell type in each diet-condition.**(F)** ATMs subtypes per gram per cohort. **(G)** Expression of key genes across ATM clusters. **(H)** *Trem2* expression in ATMs.

Immune cells were then evaluated for changes across diet conditions. ATMs increased as expected with obesity, comprising 28%, 36%, and 60% of CD45^+^ cells in mice fed ND, 8 weeks of HFD, and 14 weeks of HFD, respectively (Figure 2E, Figure S1A). Dendritic cell and monocyte populations also increased with HFD feeding, while the T cell population was highest at 8 weeks and decreased by 14 weeks of HFD feeding.

Altogether, our data capture expected WAT immune cell population dynamics in obesity progression and highlight myeloid cell accumulation in chronic obesity.

### 3.3. ATM heterogeneity spans five subtypes across early obesity

To define ATM heterogeneity, clustering was performed on ATMs from all diet-conditions (Methods Section 2.8). Five ATM subclusters were identified corresponding to resident (Mac1), proinflammatory (Mac2, Mac3), and lipid-associated (Mac4, Mac5) macrophages (Figure 2C, Methods Section 2.8).

Consistent with previous reports, resident ATMs (Mac1) expressed *Lyve1, Timd4, Mrc1/Cd206*, and *Stab1* (Figure 2C-E,G and Figure S9) [28, 29, 11].

Proinflammatory ATMs (Mac2, Mac3) were identified based on expression of genes encoding proinflammatory cytokines including *Il1b, Tnf* and *Il6* and low expression of efferocytosis markers (*Mertk, Axl, Cd163, Trem2*). Among proinflammatory ATMs, Mac2 was enriched for additional proinflammatory genes *Tnf, Il1b, Ccl2, Nlrp3* and the M2 marker *Mrc1* (Cd206). Mac3 had high expression of *Itgax/Cd11c* and antigen presentation genes (*H2-Ab1, H2-Eb1, Cd74*) and was low in *Adgre1* (F4/80), suggesting an antigen presenting ATM similar to [30]. Importantly, Mac3 was low in ATDC markers including *Zbtb46, Clec9a*, and *Cd24a* (Figure S10) [11]. Taken together, these data indicate the presence of proinflammatory macrophages that participate in monocyte recruitment and activation of T cells.

Finally, Mac4 and Mac5 ATMs emerged with HFD feeding and expressed genes consistent with lipid-associated macrophages (LAM) including *Trem2, Cd9*, and *Gpnmb* (Figure 2G) [10]. Despite transcriptional similarities, Mac4 and Mac5 differed in magnitude of LAM marker expression (Figure S8, Figure 2G). Overall, these data highlight an increase in ATM diversity with HFD feeding.

### 3.4. Lipid-associated ATMs overtake proinflammatory ATMs in chronic obesity

Next, we examined ATM phenotype dynamics during HFD feeding. To asses broad changes in the ATM transcriptional program, we examined expression of gene sets associated with phenotypic shifts in macrophages. ATMs showed progressively increased gene expression related to lipid metabolism, migration, catabolism, and cell death (Figure 2B), supporting altered metabolism and survival processes in response to obesity.

We found that resident ATMs maintained a stable population over the course of HFD feeding (Figure 2C-E). Proinflammatory macrophages were present in lean eWAT through 8w of HFD feeding but decreased substantially after 14w of HFD feeding (Figure 2C-E). In contrast, LAMs emerged with HFD feeding and continued to accumulate in chronic obesity (Figure 2C-E).

Given that other immune cells also have imbalanced subtypes in obesity and to provide additional context for ATM phenotypes during the time course, we further analyzed the single cell data for subtypes of T cells, monocytes, and dendritic cells. Known subtypes that change in adipose tissue with obesity were identified including decreased regulatory T cells and increased conventional T cells and type 2 conventional dendritic cells (Figure S7) [10, 11].

Taken together, these data show that while proinflammatory ATMs increase during adipose tissue hypertrophy, LAMs become the most prominent ATM subtype in chronic obesity.

### 3.5. LAM subtypes form a monocytic lineage

We observed that between Trem2^+^ LAMs, Mac4 outnumbered Mac5 at 8w (Figure 2D, F), but Mac5 were higher at 14w of HFD feeding (Figure 2D, F). Since LAMs are reported to be monocyte-derived [10], we hypothesized that cells in the Mac4 cluster were in transition along a monocyte-LAM lineage. Examining DE genes, 287 distinguished Mac4 and monocytes, while 834 distinguished Mac5 and monocytes (Figure S5), suggesting increasing divergence across monocytes, Mac4, and Mac5. We then queried monocytes, Mac4, and Mac5 for expression of genes related to monocyte differentiation and macrophage maturity. The monocytes markers *Cx3cr1* and *Ly6c2* were decreased in the Mac4 cluster, but were consistently higher in Mac4 compared to Mac5 (Figure 3A). Cells the Mac4 cluster also showed intermediate expression of LAM marker genes *Lgals3, Trem2* and *Ctsl* (Figure 3B). Mac4 also expressed *Ms4a7*, a marker of monocyte-macrophage differentiation, more highly than both monocytes and Mac5 [31].

**Figure 3:**
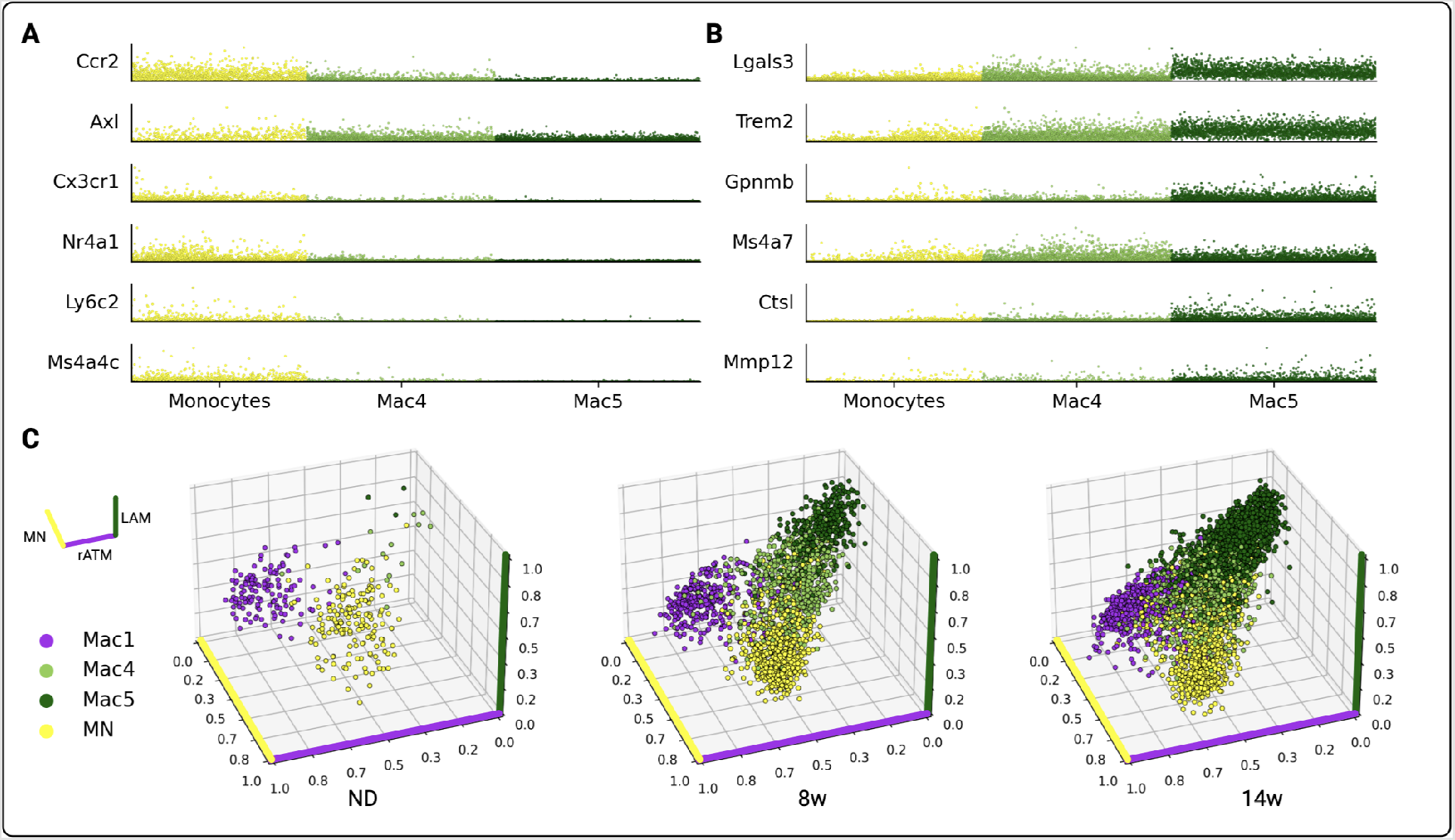
Emergence of the LAM phenotype. **(A)** Normalized expression of monocyte marker genes for key myeloid cell types. **(B)** Normalized expression of LAM marker genes for key myeloid cell types. **(C)** Three-dimensional profiling of monocytes, resident ATMs (Mac1), and LAMs (Mac4/Mac5). Cell position represents simultaneous correlation with gene expression signatures derived from monocytes (MN, yellow axis), resident ATMs (rATM, purple axis), and LAMs (green axis).

To further examine the hypothesis that Mac4 cells are pre-LAMs, we correlated them with resident ATMs (Mac1 in ND), monocytes, and chronic obesity LAMs (Mac5 in 14w). We found that Mac4 cells have intermediate correlation with the LAM and monocyte signatures, but low correlation with the resident ATM signature (Figure 3C).

Taken together, our data support that Mac4 cells are recently differentiated macrophages that are in process of acquiring the LAM phenotype.

### 3.6. Spatial transcriptomics captures LAM dynamics in obesity

The spatial context of ATM reprogramming within WAT remains poorly understood. Thus, to establish the spatial dynamics of LAM emergence with obesity, we performed spatial transcriptomics (Methods 2.5) on eWAT sampled from mice fed ND or fed a HFD for 8 or 14 weeks. We analyzed a total of 7,424 tissue capture spots across diet conditions.

Immune cell transcriptome profiles were mapped onto tissue-specific locations using conditional autoregressive-based deconvolution (CARD) (Methods 2.10) [23, 32]. We found strong emergence of the LAM phenotype across tissue spots in chronic obesity, consistent with our single cell data (Figures 4A-B, 5B, Figures S12A-B). Monocytes also increased in spatial transcriptomics data in early obesity (Figures 4A-B, 5B, Figures S12A-B). While pre-LAM spots were highest in early obesity, LAM spots were highest in chronic obesity (Figure S12B). Further, pre-LAMs and LAMs were highly spatially correlated at 8w (*r* = 0.6) but not at 14w (*r* = 0.2) (Figure S14), suggesting that LAM dynamics are spatially coordinated. Taken together, these results support LAM accumulation in WAT via differentiation from circulating monocytes.

**Figure 4:**
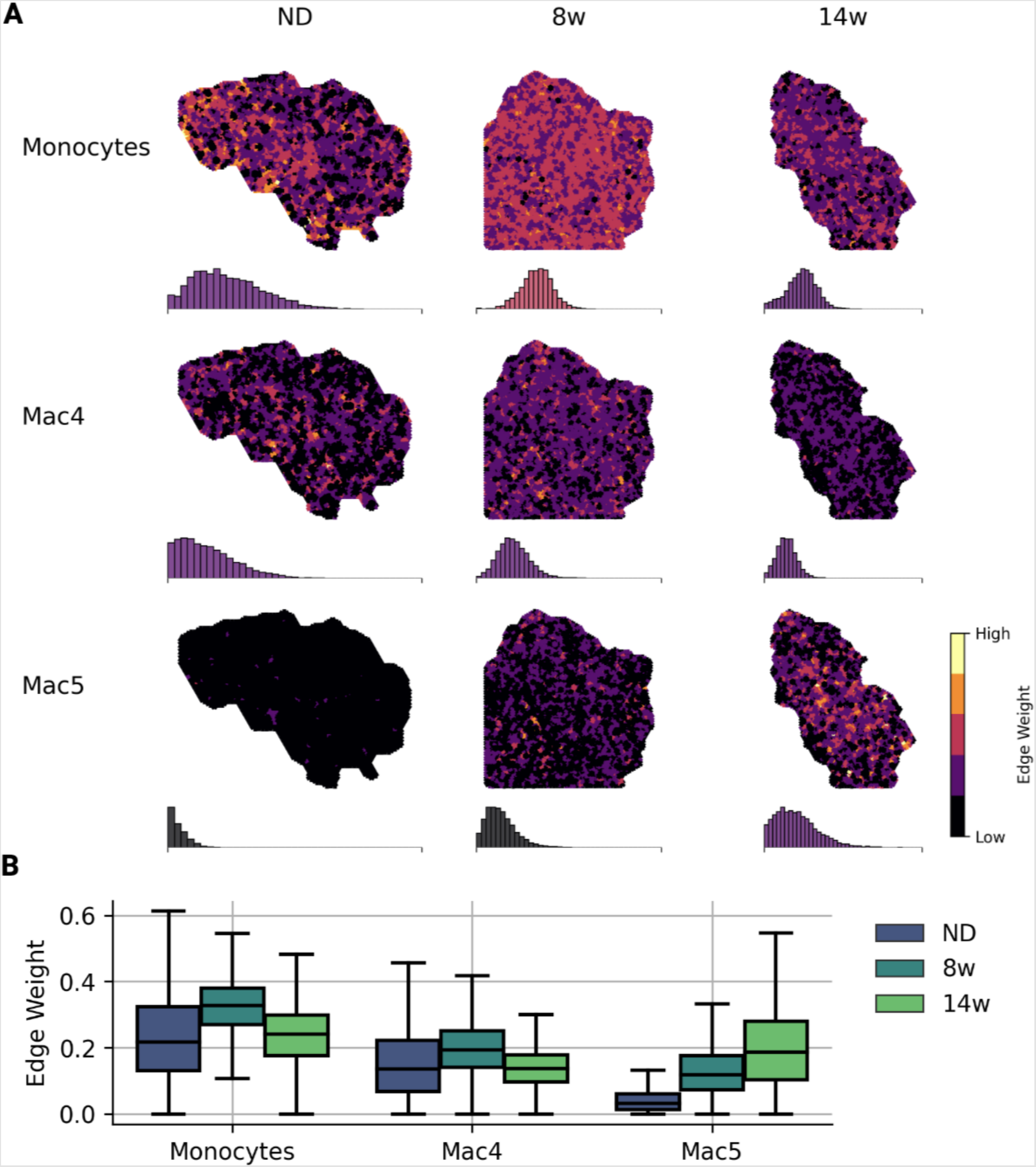
Spatial patterning of the monocyte-LAM lineage. **(A)** Spatial patterning of monocytes, pre-LAMs (Mac4) and LAMs (Mac5) over the course of HFD feeding. Edge weights are the harmonic mean of CARD proportions for neighboring capture spots. Histograms show the distribution of edge weights for the whole tissue section and are colored according to the mean edge weight on the same color scale. **(B)** Edge weight distribution by cell type and diet condition.

### 3.7. LAM networks are hubs of cell death

LAMs are associated with development of ‘crown-like structures’ (CLS), which are in turn correlated with development of insulin resistance [33, 34, 14]. CLS are well-studied [9, 35], though a spatio-temporal understanding of the drivers of CLS formation is lacking. We observed CLS as early as 8w, which prompted us to characterize the transcript patterns associated with early CLS formation. We developed cell type-specific network models based on spatial gene expression patterns and used the models to understand the dynamics of adipose tissue organization in obesity (Figure 5A, Methods 2.13).

**Figure 5:**
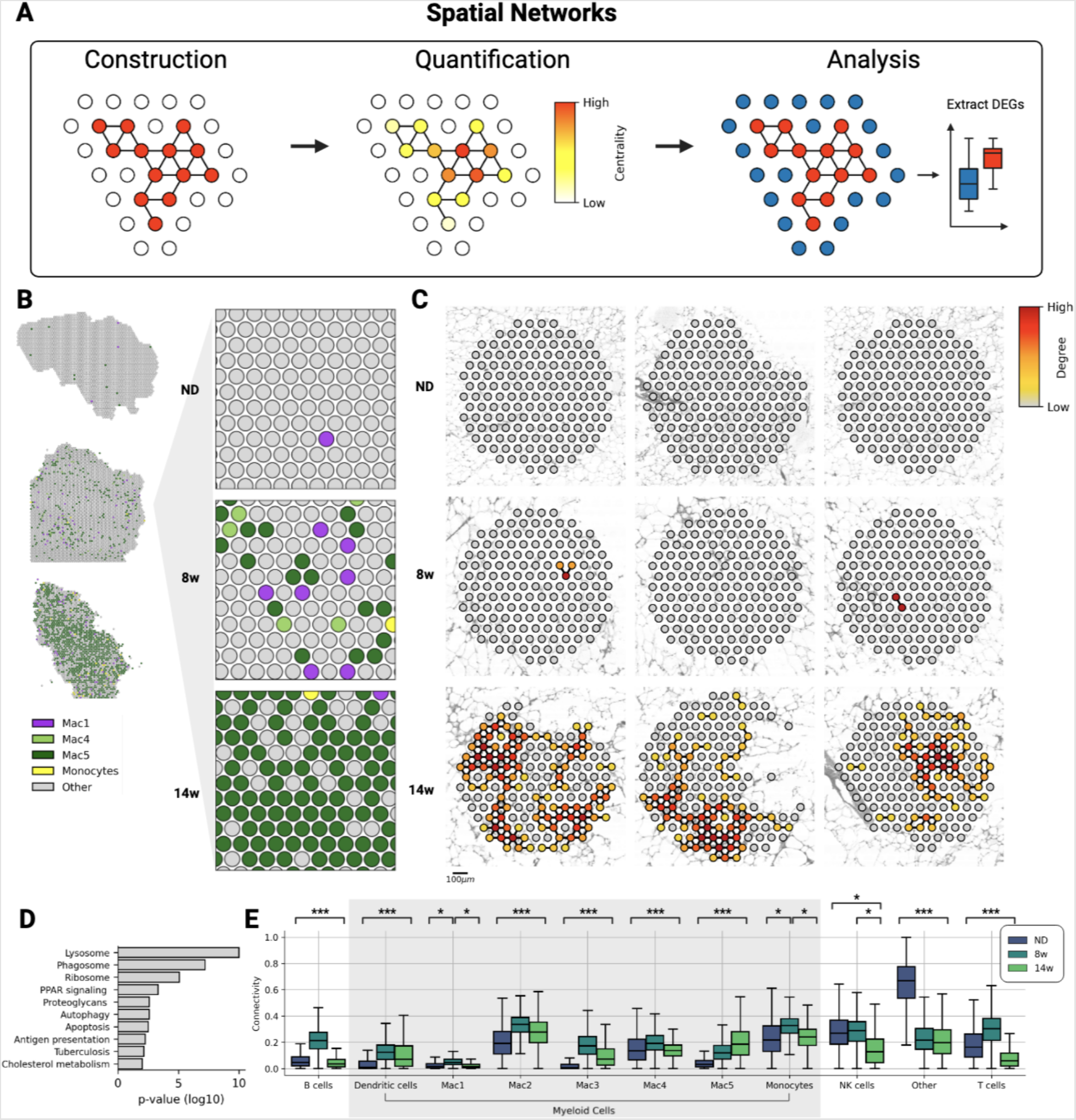
LAM networks and Hubs of Cell Death. **(A)** Workflow schematic. Network models are defined based on properties of neighboring tissue-spots. Analysis of network structure reveals principals of tissue organization. Differential expression analysis may be used to characterize the transcriptional signature of niches. **(B)** CARD-predicted cell type proportions for myeloid cell types over the course of HFD feeding. **(C)** Nine randomly sampled 150-node networks based on LAM signature (Mac5) over time. **(D)** Top 10 KEGG pathways for differentially expressed genes from LAM networks at 8 weeks and 14 weeks, compared to neighboring spatial capture spots. **(E)** Connectivity of tissue-wide networks for all immune cell types over time. Connectivity is the distribution of network edge weights, defined as harmonic mean of CARD predicted proportions between neighboring spots. Three asterisks denote that comparison between each time point (ND vs. 8w, 8w vs. 14w and ND vs. 14w) was significant (*α* = 0.05); a single asterisk denotes that the specific comparison was significant (*α* = 0.05).

Network models represent local tissue regions where a given cell type is highly localized. In the models, nodes represent tissue capture spots and edges represent interactions between adjacent nodes. Edges were defined by the harmonic mean of CARD-predicted proportions between all adjacent pairs of nodes for a given cell type (Methods 2.13). The structural properties of the cell type networks were quantified using graph-theoretic measures, which in turn revealed properties of tissue organization (Figure 5A, Methods 2.13) [26].

Network models showed higher local concentrations of adaptive immune cells (B cells, T cells) in week 8 than in lean tissue or week 14, which coincided with the emergence of proinflammatory ATMs (Figure 5E). In addition, proinflammatory Mac3 had high spatial correlation with T cells at 8w (*r* = 0.6) (Figure S14). These results suggest T cell activation, which is supported bythe emergence of T conv at 8w (Figure S7).

In contrast, local LAM concentrations increased monotonically over the course of HFD feeding, further supporting that ATM reprogramming toward the LAM phenotype is spatially coordinated. To further investigate LAM spatial patterning, we randomly sampled tissue spots from all three diet conditions and constructed 150-node networks around the sampled spot (Figure 5C). As expected, high local LAM concentrations were absent in lean tissue (Figure 5C, E). With HFD feeding, LAM concentration increased (Figure 5C, Figure S16). We then performed differential expression analysis between regions of high and low LAM concentrations and found that regions of high LAM concentrations were enriched in genes related to phagocytosis, autophagy, and cell death including *Ctsl, Ctss, Lamp1, Ctsd*, and *Ctsb* (Figure 5D). Altogether, these results identify spatially coordinated accumulation of LAMs that are engaged in clearance of excess lipids and dead adipocytes.

### 3.8. LAM networks map onto histologically identified CLS

CLS are defined by an accumulation of fibrotic and necrotic material from dead or dying adipocytes and ATMs [35, 9]. To determine the degree to which the LAM network was spatially aligned with CLS, we first developed an image segmentation algorithm to classify CLS regions from H&E images captured in parallel with spatial transcriptomics data (Figure 6A, Methods 2.14). The algorithm identified CLS_hi_ and CLS_mid_ regions of fibrotic and necrotic material that increased with obesity (Figure 6B). In contrast, area identified as adipocytes was largest in week 8 and decreased in week 14 (Figure 6C), which is consistent with adipocyte expansion in early obesity. We then aligned CLS regions with spatial transcriptomics data and found that significant colocalization of LAMs with CLS in both early and chronic obesity (Figure 6D). In contrast, pre-LAMs colocalized with CLS regions only in early obesity (Figure 4, Figure S12).

**Figure 6:**
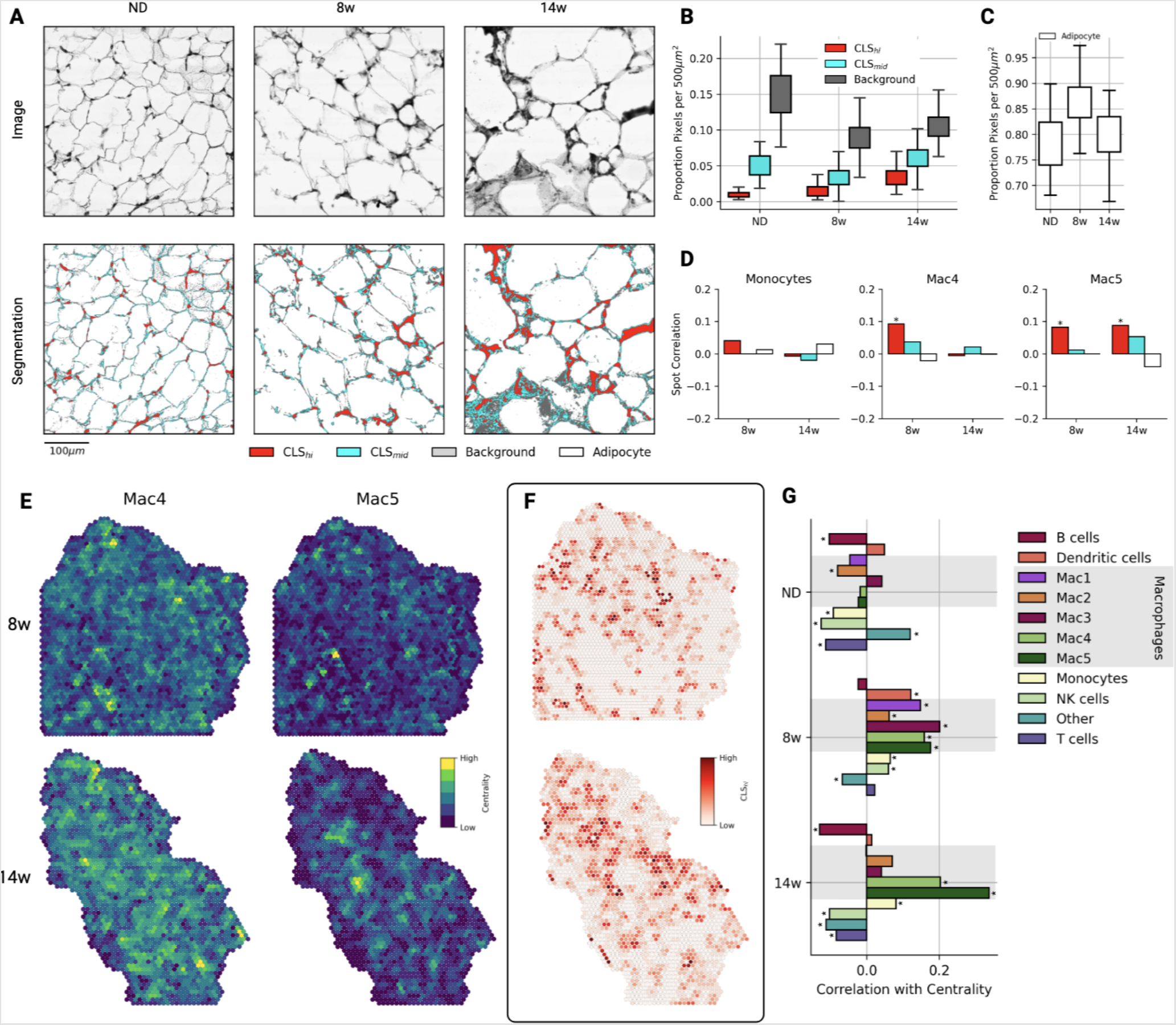
Histological Quantification of Crown-Like Structures (CLS). **(A)** H&E images captured during spatial transcriptomics library preparation (top) and segmentation results quantifying crown-like structures (bottom). **(B)** Segmentation class label proportions of 100 randomly sampled 500μm regions from each diet condition. **(C)** Adipocyte area from images regions in (B). **(D)** Spot correlation between myeloid cell type proportions and segmentation results from a 150μm region around each capture spot. Asterisks denote significant Pearson correlation values (*α* = 0.01). **(E)** Spot importance in global cell type networks (eigenvector centrality) in HFD feeding conditions. Eigenvector centrality highlights regions of densely localized cells in the tissue. **(F)** CLS_hi_ segmentation results in 150μm regions around each capture spot at 8 and 14 weeks. **(G)** Spot correlation between CLS_hi_ segmentation results and eigenvector centrality for each diet-condition, by cell type. Asterisks denote significant Pearson correlation values (*α* = 0.01).

Beyond correlation, we sought to characterize the physical organization of immune cell types within WAT and their relationship to CLS. We used eigenvector centrality, a global measure of nodal importance in a network, to quantify cell type-specific structure within the tissue [26]. We then correlated per-spot centrality for each immune cell type network with per-spot CLS prevalence (Figure 6G). We found that critical hubs of innate immune cells aligned with early CLS in week 8 (Figure 6G). Central nodes in pre-LAM and LAM networks aligned with CLS both in early and chronic obesity (Figure 6E-G). In contrast, adaptive immune cell types (B cells, T cells) exhibited negative correlation with CLS in all diet conditions.

Taken together, these results capture the dynamic, large-scale reorganization of immune cells in early obesity and the spatial concentration of LAMs in CLS regions in chronic obesity.

### 3.9. Myeloid signaling shapes nascent CLS

Given the early presence of CLS and reorganization of myeloid cell types in week 8, we sought to characterize intracellular signaling during formation of CLS. We therefore quantified spatially colocalized expression of ligand-receptor (LR) pairs throughout WAT and within the monocyte-LAM lineage.

We first cataloged tissue-wide changes in LR expression. We identified the LR pairs that increased in early obesity and chronic obesity (Figure 7A-B) and the LR pairs that decreased in early and chronic obesity ((Figure 7C-D, Methods 2.12). As expected, global LR analysis revealed increased metabolic activation (*Lrp1, Lpl, App, Apoe*), regulation of cellular migration (*Adipoq, Igf1, Thbs1, Apoe*), regulation of tissue remodeling (*Cola1, Cola2*) and regulation of immune response (*Cd36, Cd81, C3*) (Figure 7A-D, Figure S15) as predominant biological processes associated with obesity-induced WAT remodeling.

**Figure 7:**
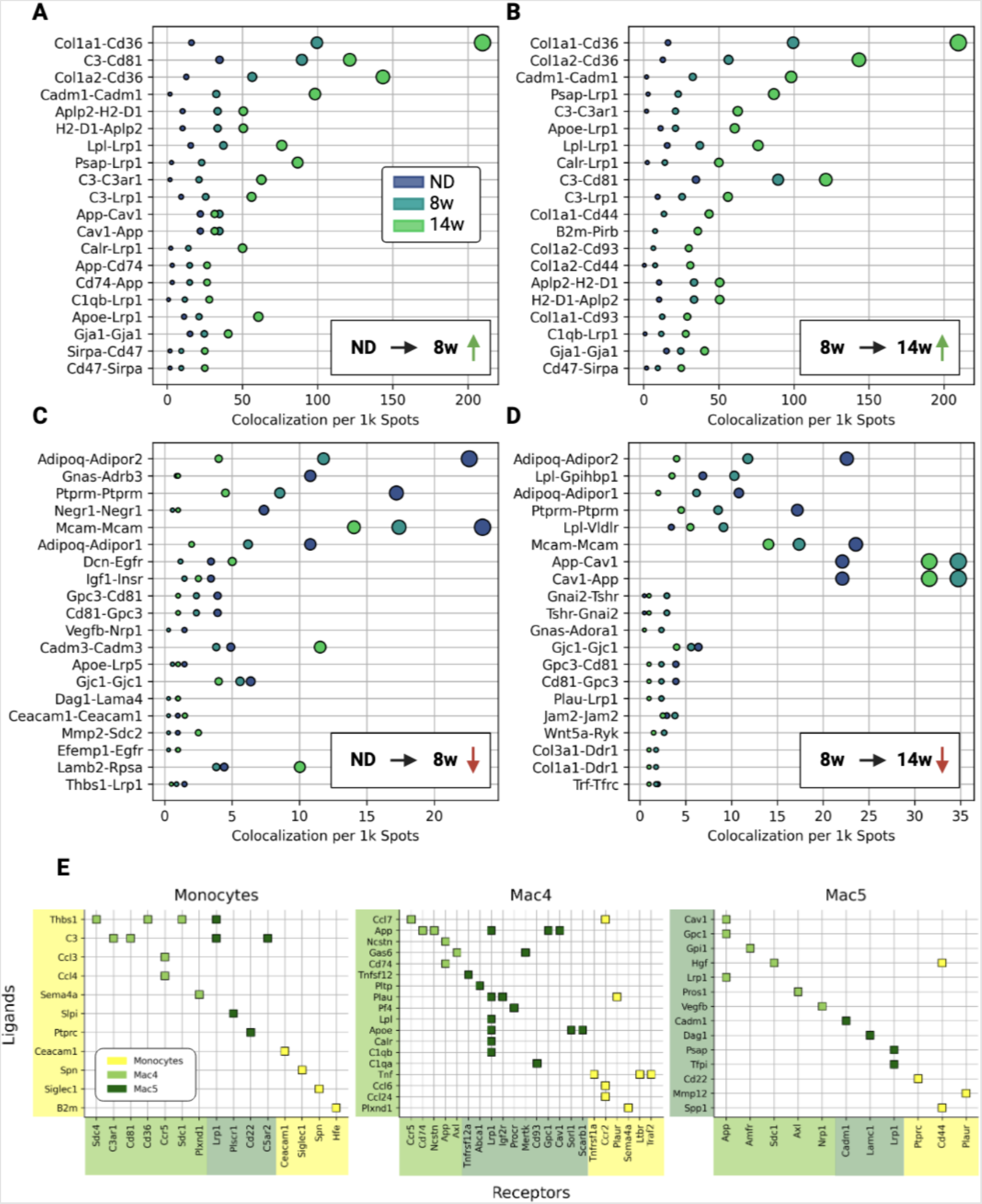
WAT ligand-receptor signaling dynamics. **(A)** Ligand-receptor (LR) pairs with most increased colocalization during the first 8 weeks of HFD feeding. Dot sizes are LR colocalization per 1k capture spots (same as x-axis) and dot colors indicate diet-condition. **(B)** LR pairs with most increased colocalization during the last 6 weeks of HFD feeding. **(C)** LR pairs with most decreased colocalization during the first 8 weeks of HFD feeding. **(D)** LR pairs with most decreased colocalization during last 6 weeks of HFD feeding. **(E)** Differential expressed myeloid LR pairs with non-zero colocalization in spatial data.

To identify the myeloid-specific signaling that may contribute to the emergence of CLS, we investigated LR pairs that were both differentially expressed in a myeloid cell subtype and colocalized with one another in the spatial transcriptomics data (Figure 7E, Methods 2.12). Pre-LAMs expressed multiple ligands for LAM receptor *Lrp1*, including *App, Plau, Lpl, Apoe, Calr* and *C1qb*. Additionally, pre-LAMs expressed ligands *App, Plau, Apoe* that had multiple receptors throughout the monocyte-LAM lineage.

Thus, we identify a novel set of signaling molecules expressed in early obesity along the monocyte-LAM lineage that may significantly influence the nascent CLS microenvironment.

## 4. DISCUSSION

Changes in mammalian adipose tissue immune cells persist even in weight loss [15, 12], highlighting the need to better understand mechanisms that promote adipose tissue dysfunction. Our study elucidates ATM phenotype dynamics in their spatial context in early and chronic obesity by combining single cell RNA-seq, spatial transcriptomics, and imaging over time.

Our work supports increased phenotypic diversity in ATMs with obesity that is consistent with other single cell work [10, 24, 35, 11]. Our data captured the dramatic increase in ATMs that were phenotypically distinct from resident ATMs in lean tissue (Figure 3H), and ATMs overall showed metabolic and catabolic activation in obesity (Figure 3A). We also show that the LAM phenotype became dominant among ATMs in chronic obesity [10, 8, 14] 3B-G). These data are consistent with other work demonstrating that ATMs acquire non-classical activation states in obesity [36, 37, 14, 24].

LAMs are reported to be anti-inflammatory, tissue-remodeling macrophages that are highly metabolically active; their transcriptional signature is associated with phagocytosis and endocytosis [13] and they have elevated expression of markers such as *Trem2, Lgals3* and *Ctsl* [10]. Our data agree with these findings and additionally identify a novel population of pre-LAMs as a closely related precursor to LAMs (Figure 3H).

Significant appearance of pre-LAMs precedes accumulation of LAMs and coincides with initial formation of CLS. Spatial analyses further support pre-LAM localization to CLS in early obesity and suggest pre-LAM signaling through *App, Apoe, Lpl*, and *Lrp1* as drivers of CLS formation.

These molecules implicate disruption of lipid processing pathways in development of tissue dysfunction. Dysregulated lipid processing is associated with oxidative and ER stress that alters cell survival and macrophage phenotype [38, 39, 40, 41], which are in turn hallmarks of disease progression in type II diabetes and neurological disorders [42, 41].

Limitations of this study include low cell numbers in our single cell data (1.2k-6.4k cells), which limits identification of rare but functionally important cell types. Although we identified multiple ATM subtypes, other immune cell subtypes were less identifiable, potentiality due to low cell numbers. Known shifts in subtypes include increased CD8^+^ T effector and CD4^+^ T_H_1 cells and decreased regulatory T cells in obesity [43, 44, 45]. In addition, spatial transcriptomics data included only one tissue section per diet condition and were relatively low depth with a median of 91-173 genes identified per capture spot. We therefore used nearby capture spots to improve cell type identification at each spot used nearby capture spots to infer cell type proportions at each capture spot [23]. Finally, data were only collected from male mice which limits comparisons based on sex.

## Conclusions

Our data revise current understanding of ATM phenotypic shifts in obesity. We identify important milestones in monocyte-LAM development and provide spatial context for myeloid signaling that is implicated in metabolic dysfunction. Our study provides novel clarity on the cell types and signaling involved in CLS formation and accumulation, including the spatial dynamics of lipid-associated macrophage development in obesity.

## Author contributions

**Cooper Stansbury:** Methodology, software, formal analysis, data curation, writing and editing the manuscript, visualization; **Gabrielle A. Dotson:** Methodology, software, formal analysis, data curation, writing and editing the manuscript, visualization; **Harrison Pugh:** methodology, formal analysis, and writing and editing the manuscript; **Alnawaz Rehemtulla:** methodology, resources, writing and editing the manuscript; **Indika Rajapakse:** methodology, resources, writing and editing the manuscript, supervision; **Lindsey A. Muir:** conceptualization, methodology, investigation, formal analysis, resources, writing and editing the manuscript, visualization, supervision.

## Acknowledgements

We thank Ingrid Bergin and Pavlina Zafirovska for technical assistance in sectioning and staining tissues, Olivia Koues and the University of Michigan Advanced Genomics Core, Evan Keller and Greg Shelley for assistance with single cell sequencing, Abigail Riesmeyer and Siva Kumar for technical assistance, and Stephen Lindsly and Can Chen for their input on data analysis. We also acknowledge BioRender, which was used to generate all figures.

## Data and resource availability

The spatial transcriptomics and single-cell RNA-seq datasets generated in this study have been deposited to the Gene Expression Omnibus (GEO) and can be accessed via accession number GSE198012.

