## Supplemental Figures and Tables for "A lipid-associated macrophage lineage rewires the spatial landscape of adipose tissue in early obesity"

### SUPPLEMENTARY MATERIAL

#### Supplementary Figures

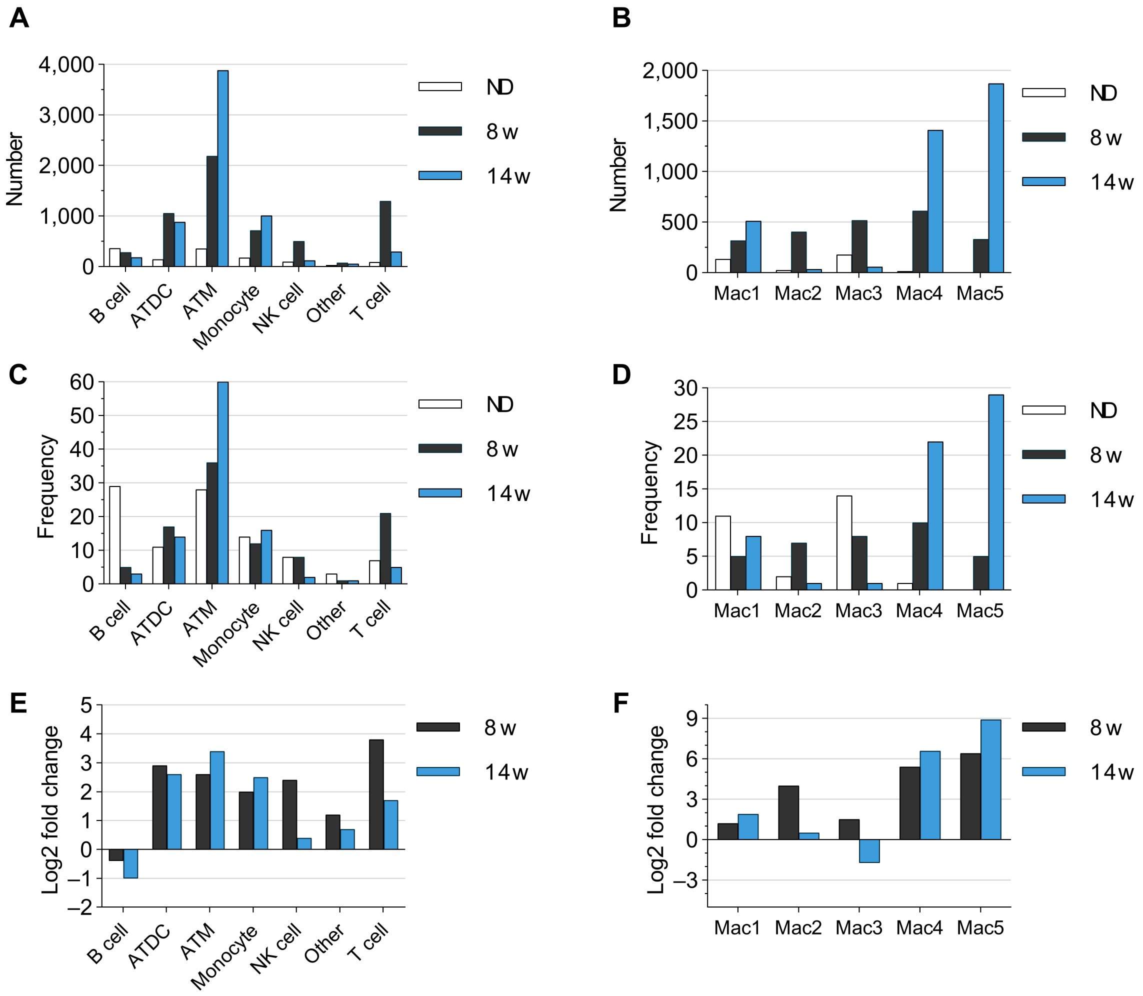


Supplementary Figure 1: **Immune cell scRNA-seq summary.** **(A)** The total number of cells of each type by diet condition. **(B)** The total number of macrophages in each subcluster by diet condition. **(C)** The frequency of each cell type by diet condition. **(D)** The frequency of each macrophage subcluster by diet condition. **(E)** The fold change (log2) of cell number over ND for all cell types. **(F)** The fold change (log2) of cell number over ND for all macrophage subtypes.

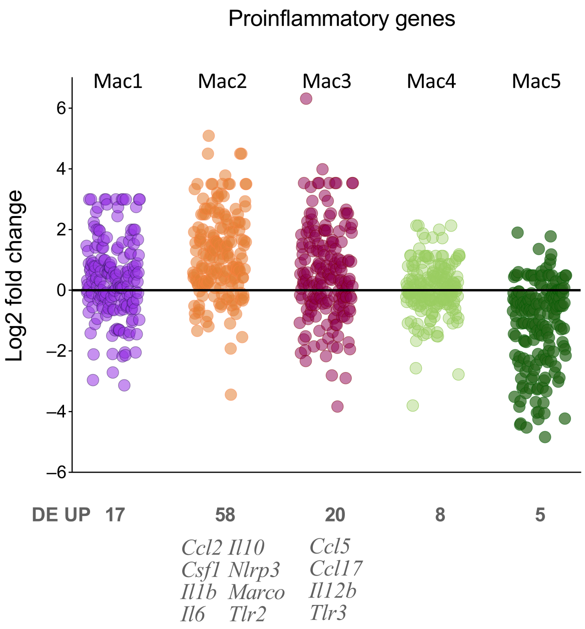


Supplementary Figure 2: **Inflammatory response-related gene expression in ATM subtypes.** Each point represents log2 fold change in expression of a gene for the given ATM subtype compared to all other subtypes at 8w of HFD feeding. Genes were included in the plot if they were DE for at least one ATM subtype (185 of 197 genes in the mouse gene set). The number of DEGs that were increased (DE UP) is given for each ATM subtype. Select DE UP genes are shown for Mac2 and Mac3. DE genes among ATM subtypes were determined using 10X Genomics Loupe Browser v6.4.0 using the "Locally Distinguishing" comparison.

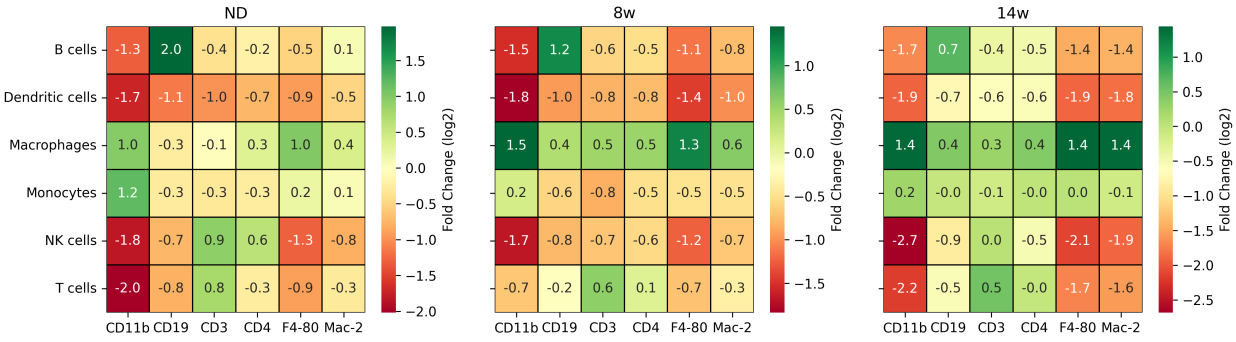


Supplementary Figure 3: **Feature barcoding summary.** Protein expression fold changes with respect to cell type annotations across all three diet conditions.

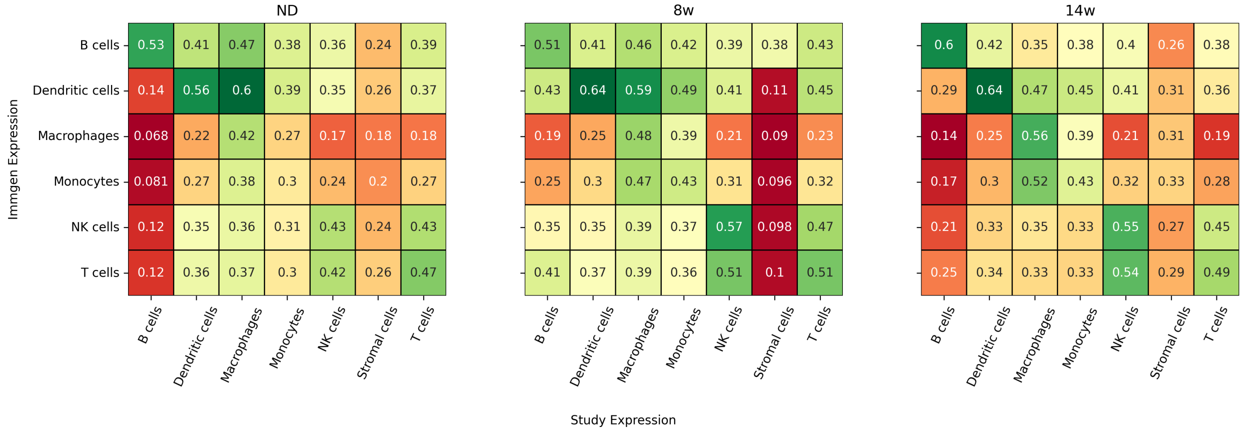


Supplementary Figure 4: **Correlation of cell type annotations with ImmGen expression profiles.** Correlation of normalized expression from each cell type at each time point with ImmGen profiles (GSE122597, GSE124829, GSE75202, GSE15907, GSE75203, GSE122108, GSE37448, GSE109125, and GSE110549) for the same cell types [46].

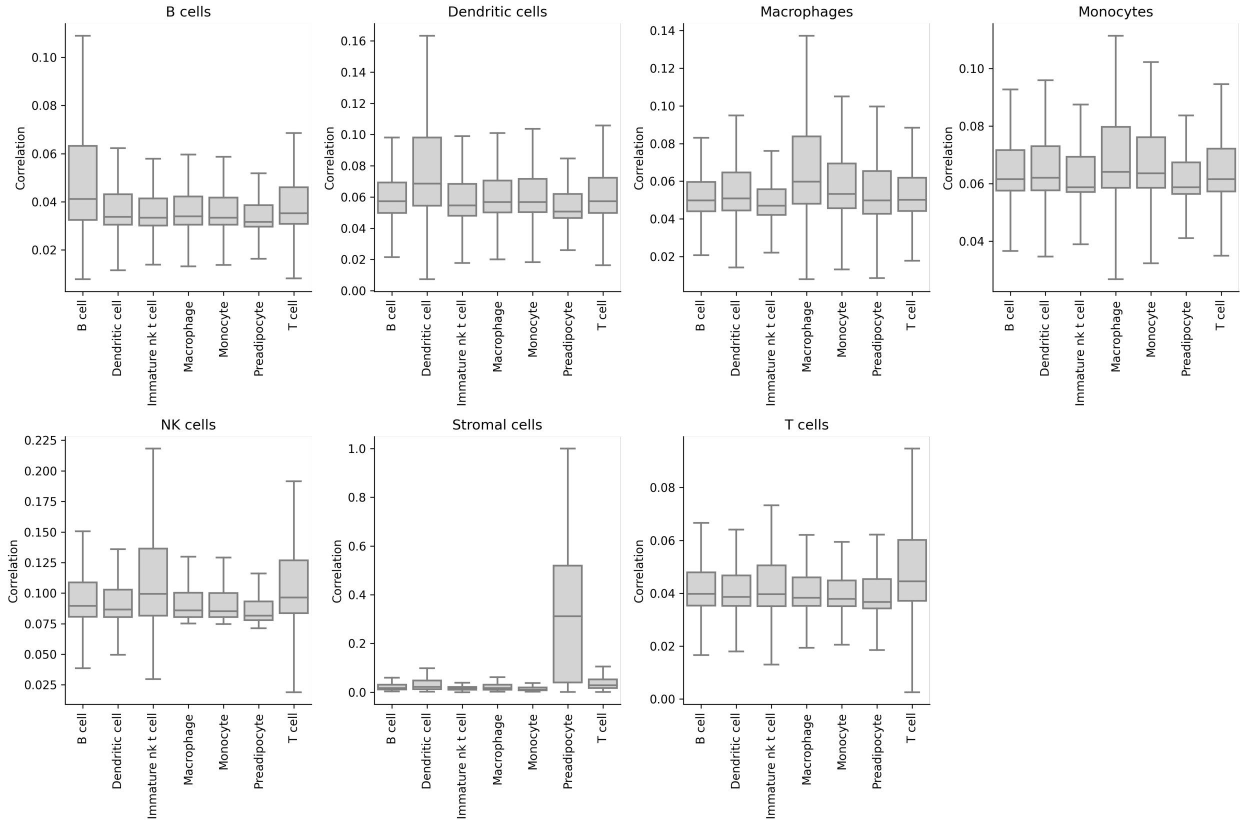


Supplementary Figure 5: **Cell type annotations align with murine eWAT scRNA-seq profiles.** Normalized expression of genes in the top 50-percentile of variance in our data was correlated with aggregated expression profiles from [8].

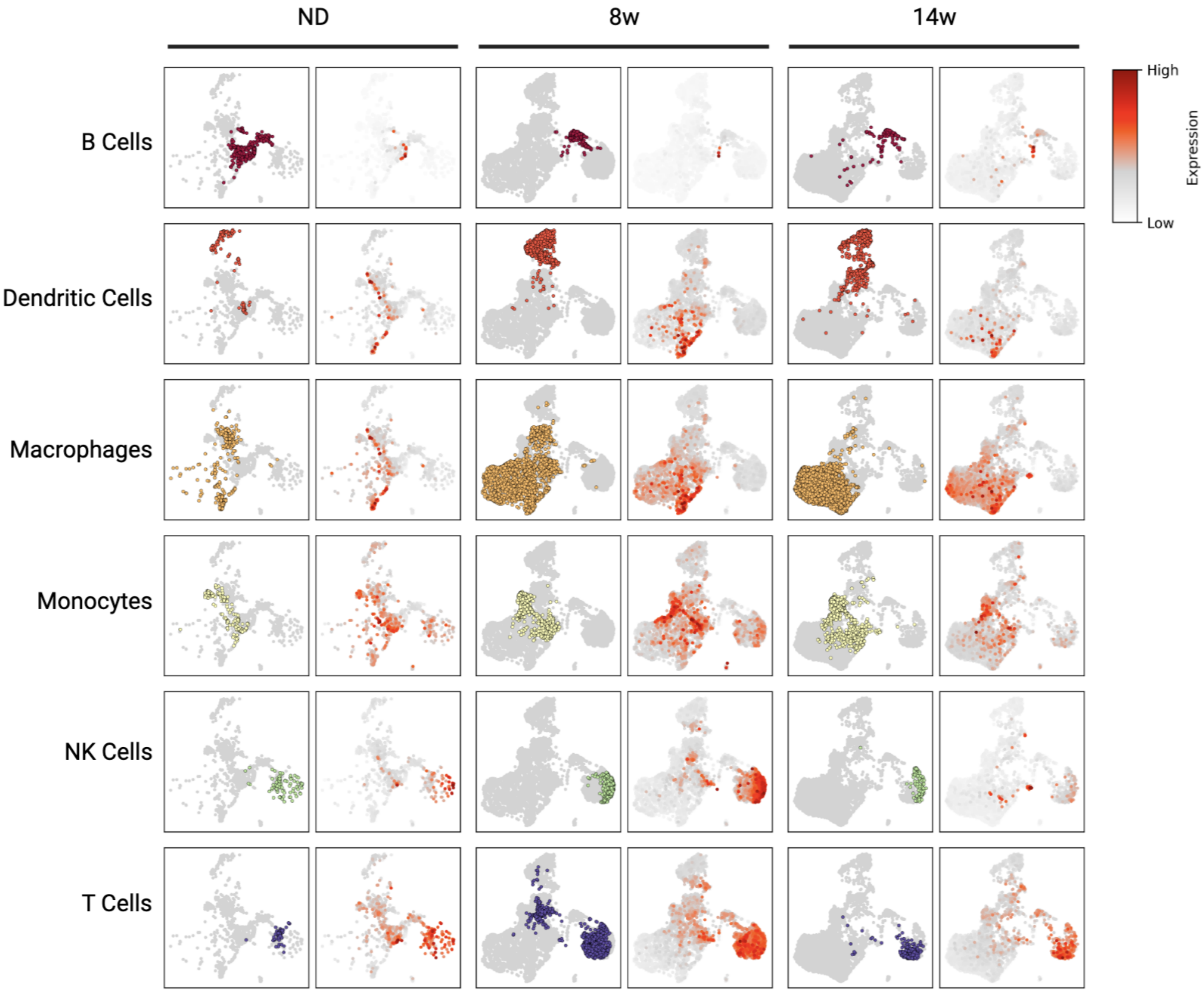


Supplementary Figure 6: **PanglaoDB marker gene expression by cell type.** Square-root mean expression of mouse marker genes from PanglaoDB (UI < 0.025) [22].

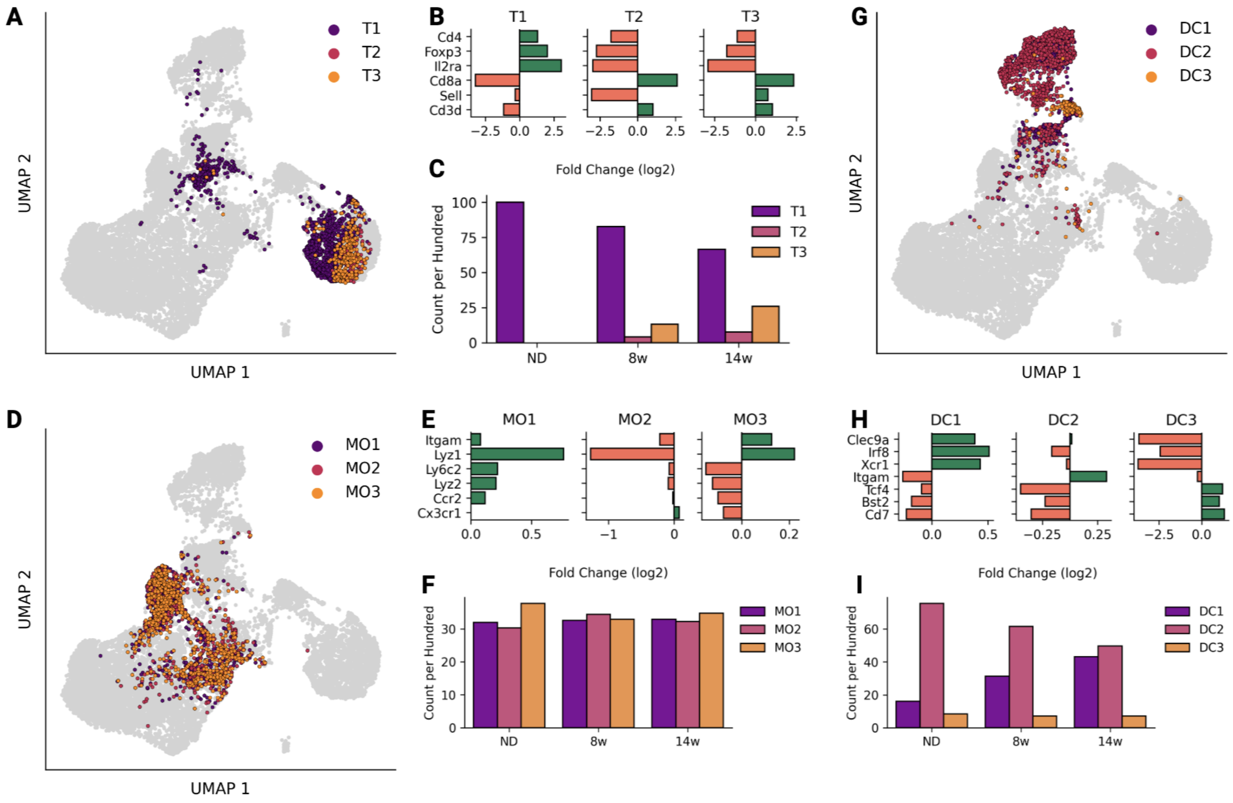


Supplementary Figure 7: **Adipose tissue T cell, monocyte, and dendritic cell subtypes in obesity by scRNA-seq.** **(A-C)** T cell subclusters included regulatory T cells (T1), conventional T cells (T2), and all other T cells (T3). **(A)** UMAP of T cell subclusters. **(B)** Differentially expressed T cell subtype markers shown as log2 fold change of each population versus all others. **(C)** T cell subtype quantity per diet condition, shown as count per hundred (proportion of parent). **(D-F)** Monocyte subclusters included inflammatory (MO1, MO3) and patrolling (MO2) subtypes. **(D)** UMAP of monocyte subclusters.**(E)** Differentially expressed monocyte subtype markers shown as log2 fold change of each population versus all others. **(F)** Monocyte subtype quantity per diet condition, shown as count per hundred (proportion of parent). **(G-I)** Dendritic cell subclusters included classical (DC1, DC2) and plasmacytoid (DC3) subtypes. **(G)** UMAP of dendritic cell subclusters. **(H)** Differentially expressed dendritic cell subtype markers shown as log2 fold change of each population versus all others. **(I)** Dendritic cell subtype quantity per diet condition, shown as count per hundred (proportion of parent). DE genes were determined using Wilcoxon rank sum tests.

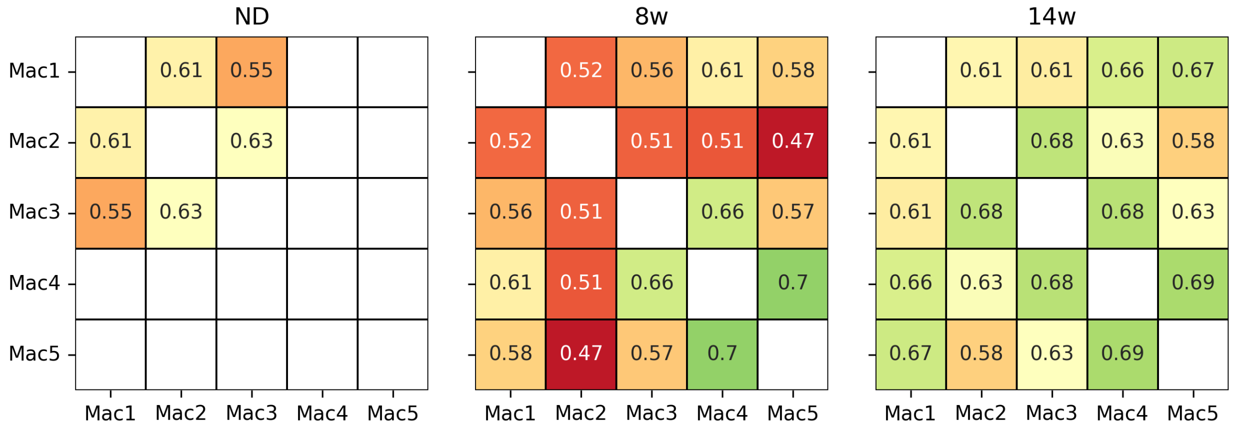


Supplementary Figure 8: **Macrophage subtype genomics correlation over time.** Correlation between macrophage subtypes over PCA embedding (95% explained variance) generated from genes in at least 5% of all macrophages. Correlations are only shown when the subtype has more than 50 cells in the given diet-condition.

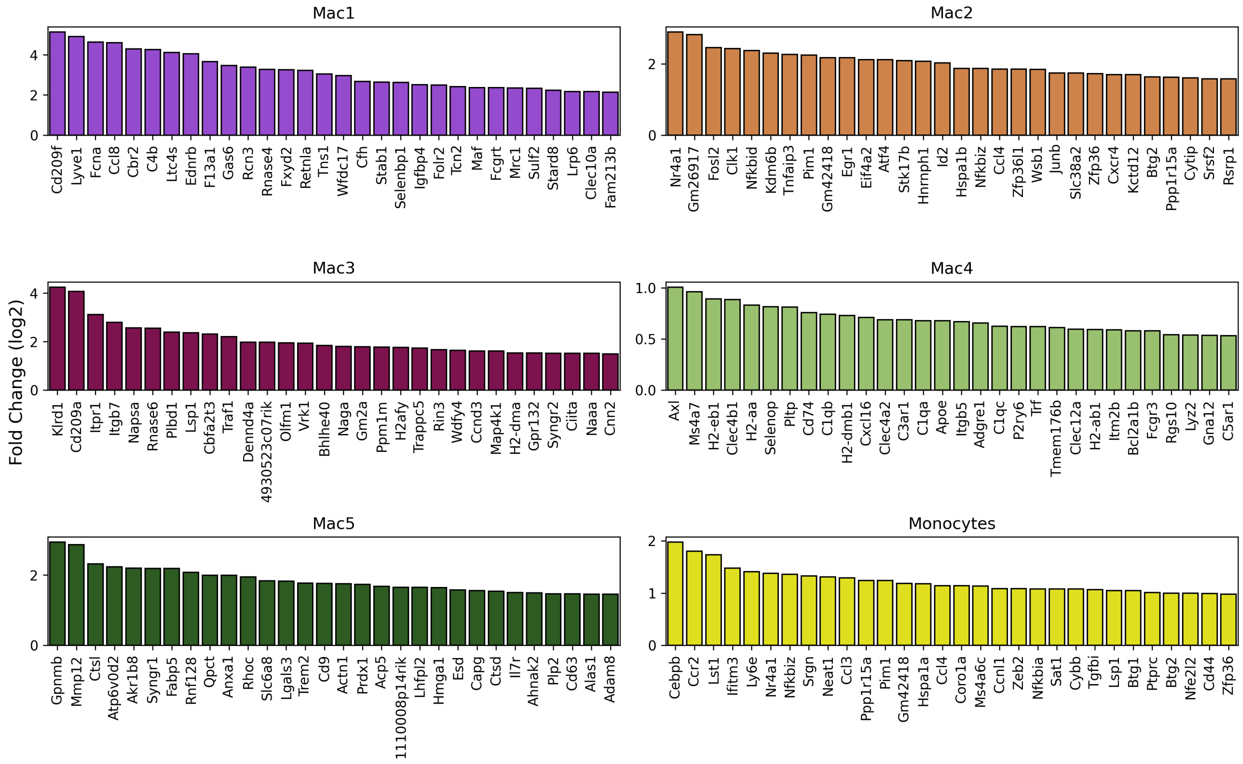


Supplementary Figure 9: **Differential expressed macrophage subtype genes.** Top differentially expressed genes from non-parametric Wilcoxon rank-sum tests adjusted using Bonferroni correction ($\alpha = 0.05$). Genes are expressed in at least 50% of the cell type, arranged by descending fold change (log2).

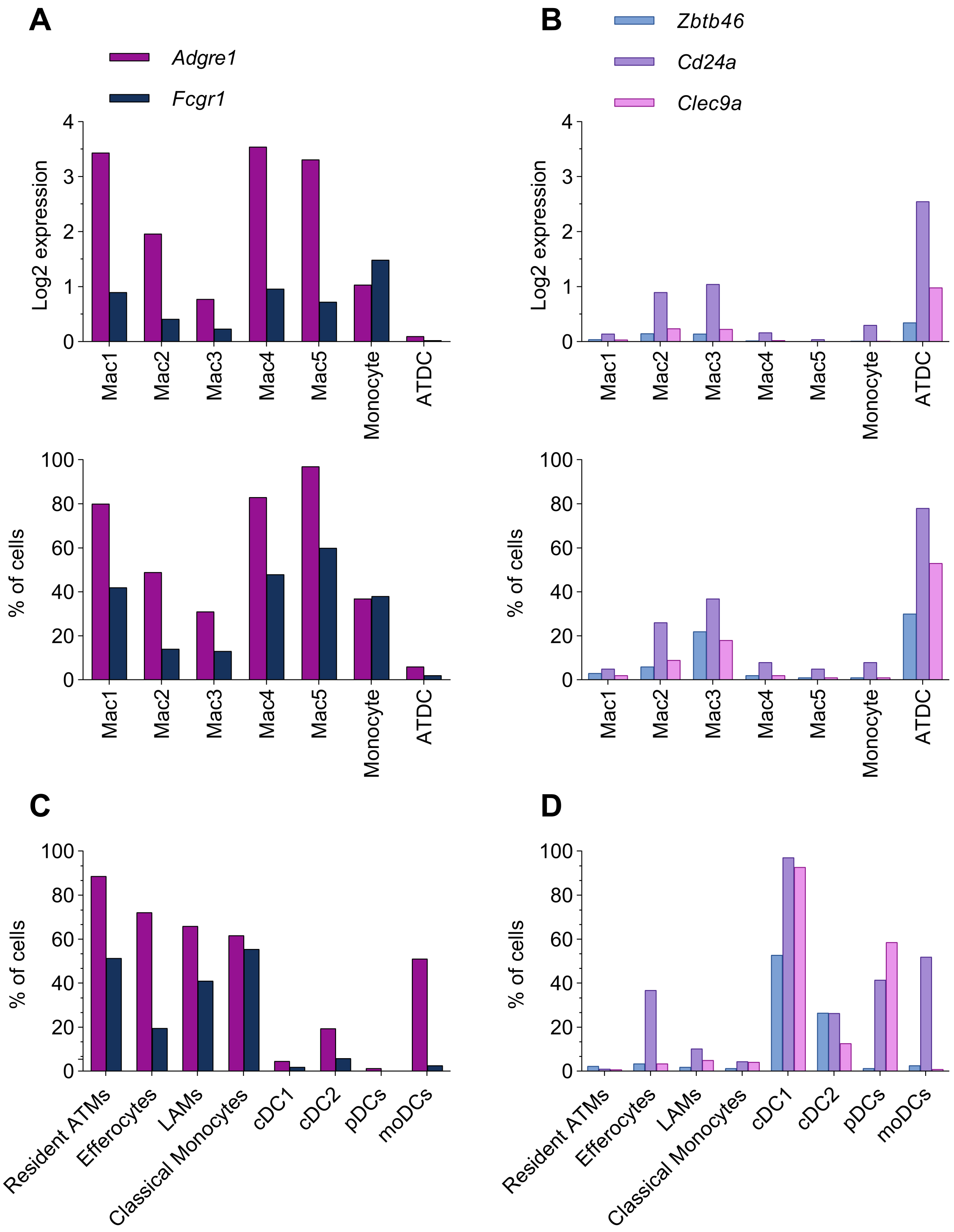


Supplementary Figure 10: **Macrophage and dendritic cell marker genes.** **(A)** Expression of macrophage marker genes. **(B)** Expression of dendritic cell marker genes.**(C-D)** Expression of macrophage (C) and dendritic cell (D) marker genes in macrophage, monocyte, and dendritic cell subsets from [11]. ATDC: adipose tissue dendritic cells.

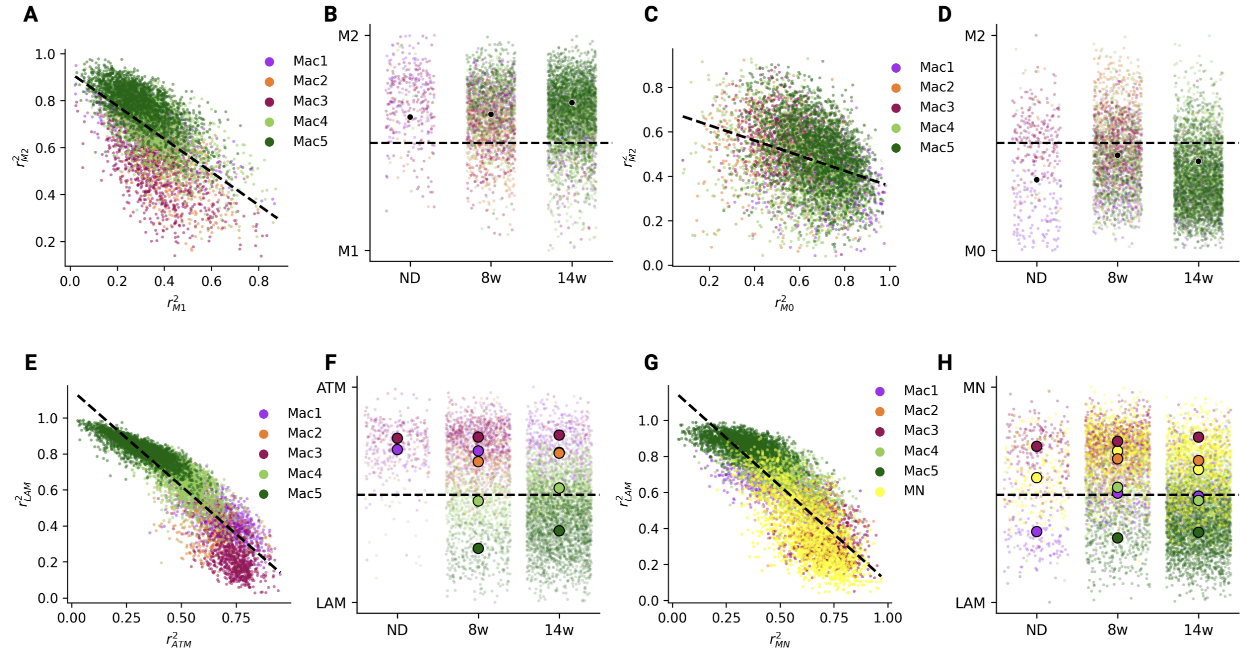


Supplementary Figure 11: **Macrophage polarization states.** **(A)** Correlation of macrophages to M1 and M2 macrophages from GSE117176 based on the 100 DEGs sorted by fold change (log2). **(B)** Macrophage polarization state determined by distance from the upper-left along the line of best-fit (A). **(C)** Correlation of macrophages to M2 and M0 macrophages from GSE117176 based on the 100 DEGs based on fold change (log2). Macrophage polarization state determined by distance from the upper-left along the line of best-fit (C). **(E)** Correlation of macrophages to resident macrophages (Mac1, ND) and LAM macrophages (Mac5, 14w) based on the 100 DEGs sorted by fold change (log2). **(F)** Macrophage polarization state determined by distance from the upper-left along the line of best-fit (E). **(G)** Correlation of macrophages to monocytes and LAM macrophages (Mac5, 14w) based on the 100 DEGs sorted by fold change (log2). **(H)** Macrophage polarization state determined by distance from the upper-left along the line of best-fit (G).

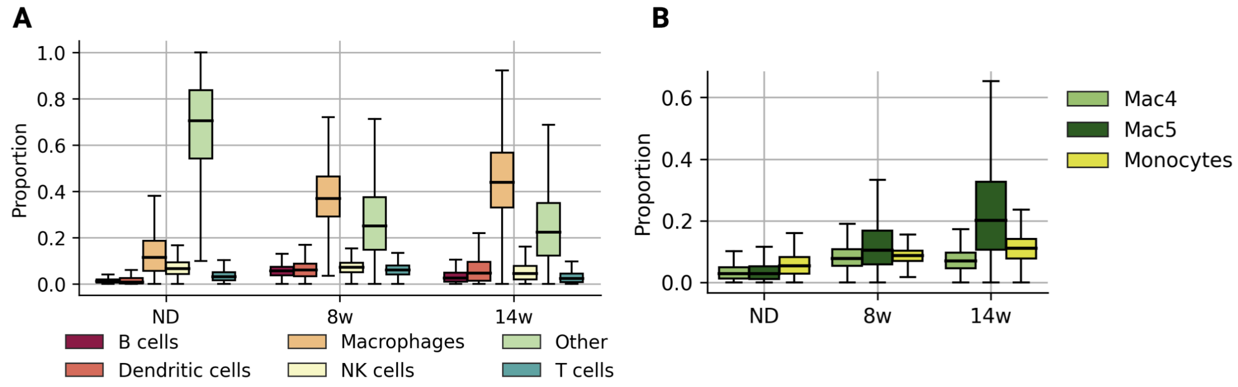


Supplementary Figure 12: **Tissue-capture spot cell-type proportions over time.** **(A)** CARD-estimated immune-cell type composition of each capture spot at each diet-condition. **(B)** CARD-estimated monocyte-LAM lineage cell-type composition at each diet condition.

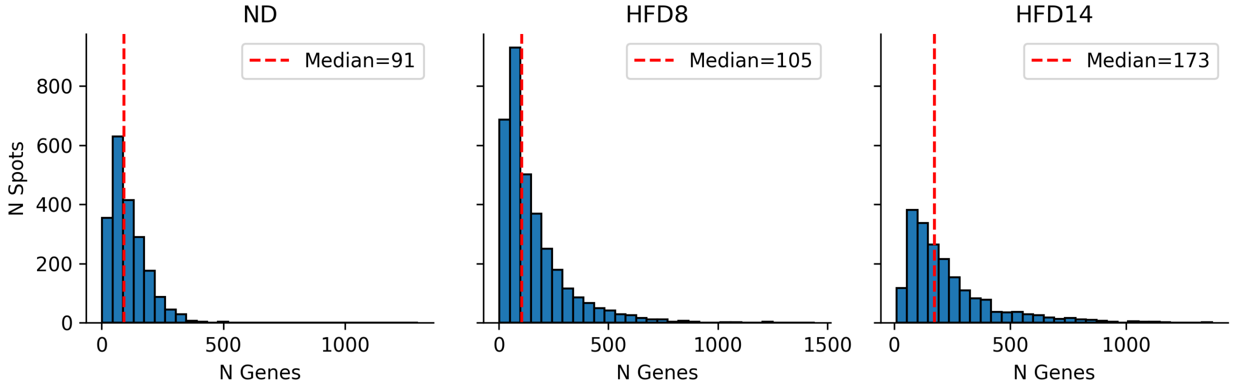


Supplementary Figure 13: **Spatial transcriptomics read depth summary.** Distribution of detected transcripts per spot in spatial transcriptomics data at eadh diet condition.

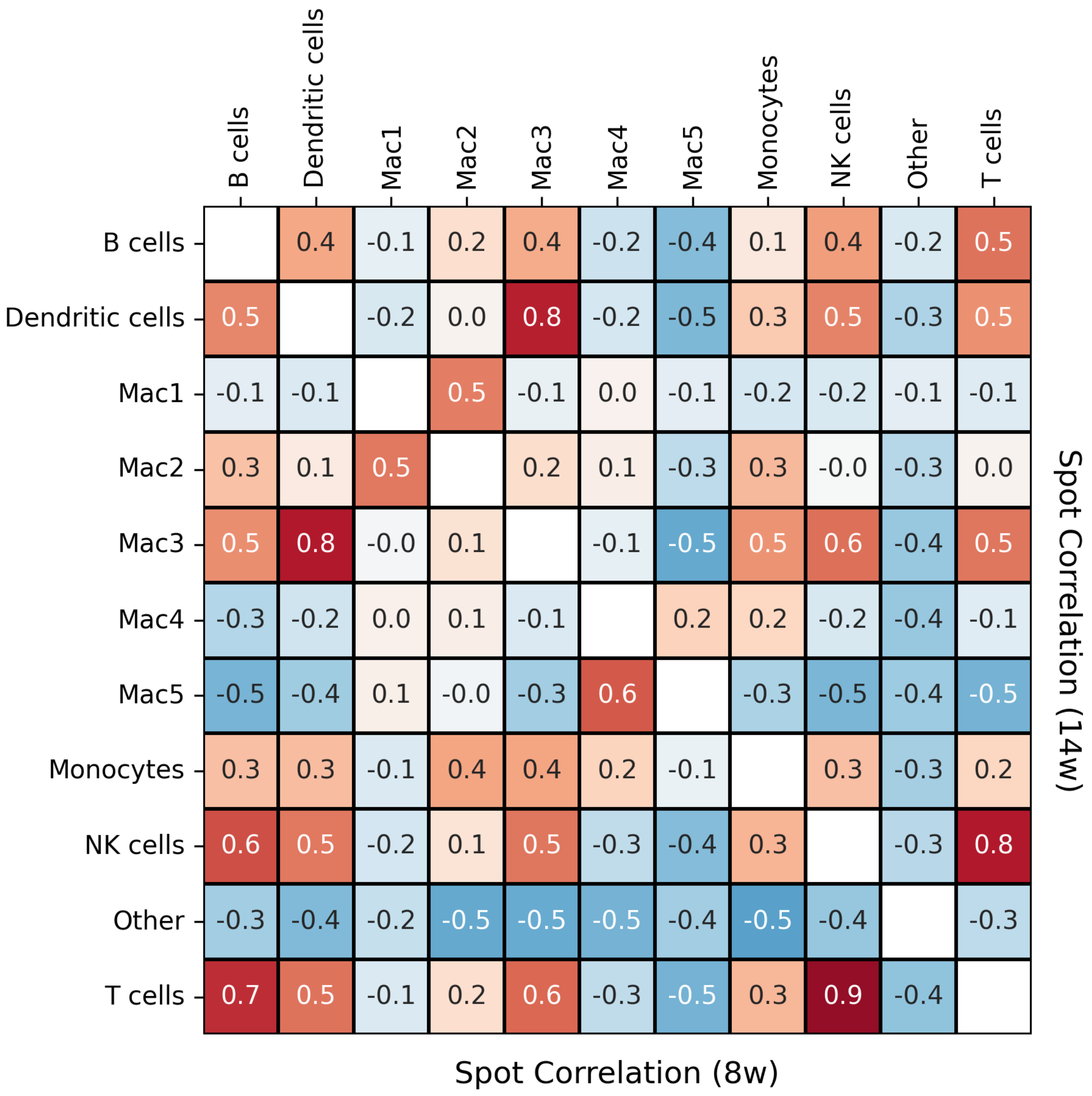


Supplementary Figure 14: **Immune-cell spot correlations.** Correlation of predicted CARD proportions of immune-cell types over all tissue-capture spots at 8 weeks (below diagonal) and 14 weeks (above diagonal).

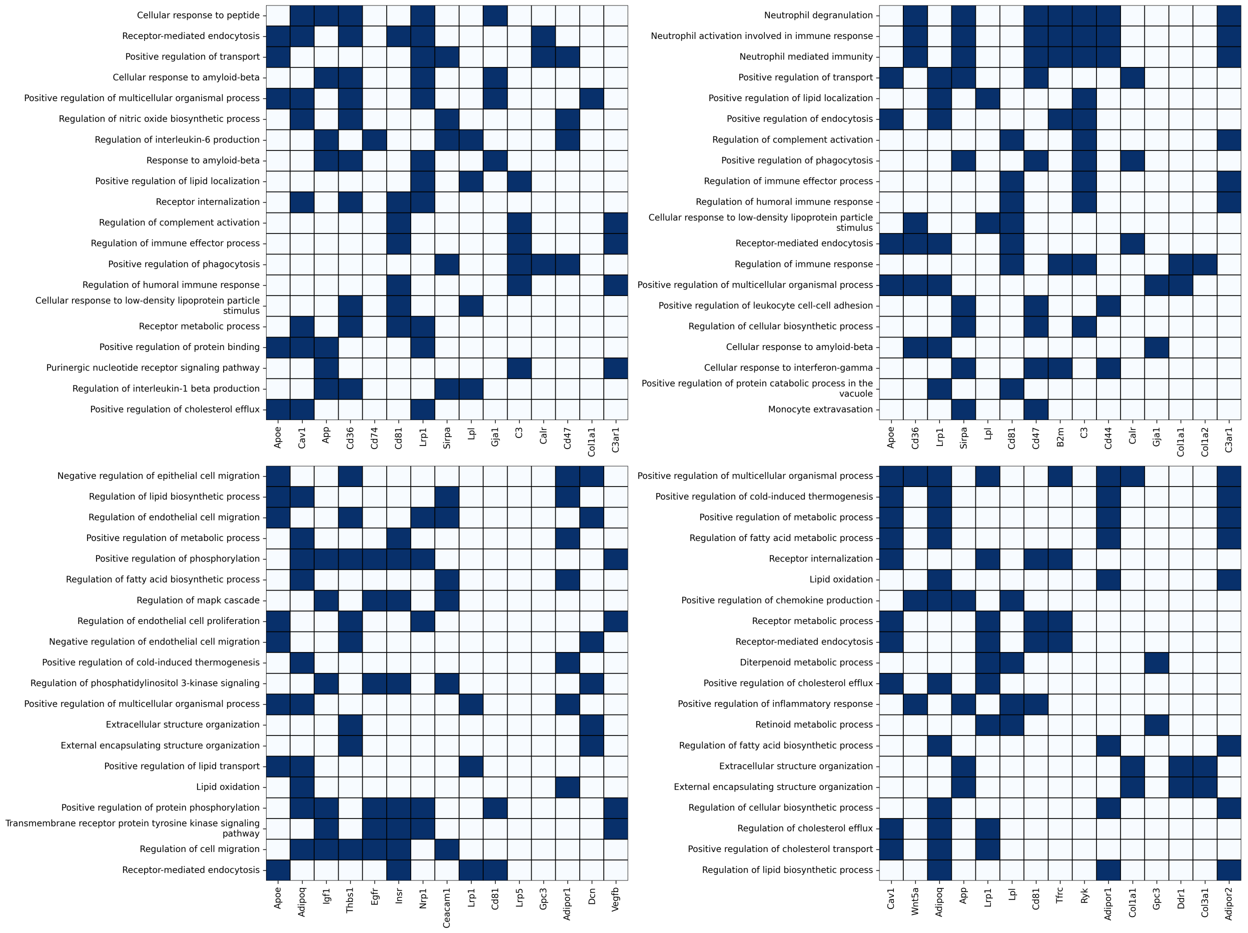


Supplementary Figure 15: **Ligand-Receptor colocalization reveals key biological processes during HFD feeding.** Enriched Gene Ontology biological processes for colocalized ligand-receptor (LR) pairs. LR pairs are divided into four groups: (upper left) increased in the first 8 weeks, (upper right) increased between 8w and 14w, (lower left) decreased in the first 8 weeks and (lower right) decreases 8w and 14.

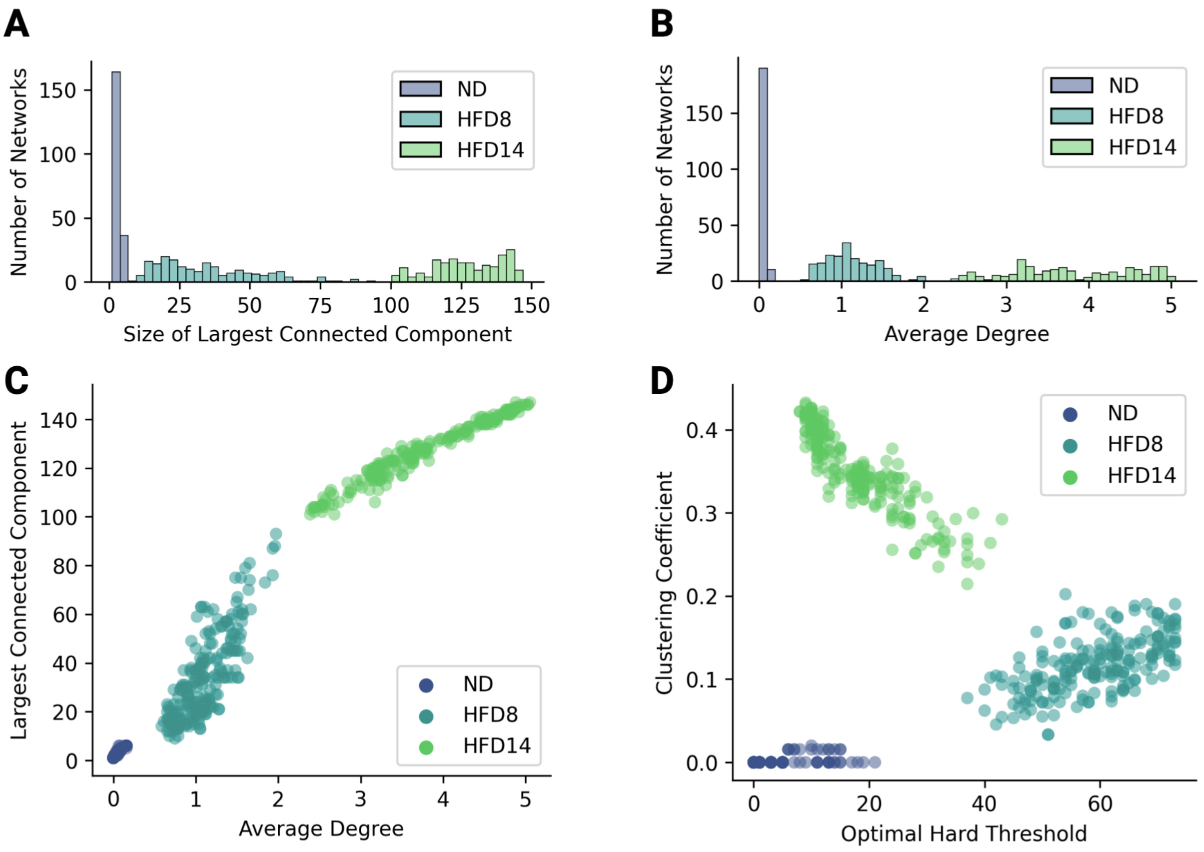


Supplementary Figure 16: **LAM networks show increased connectivity with HFD feeding.** **(A)** Size of largest fully connected component for LAM networks edge-thresholded at 0.015 for each diet condition. A connected component is defined by the set of nodes which can be traversed by travelling on defined edges. The largest connected component of a graph is the largest set of nodes connected by at least 1 edge. **(B)** The mean number of edges for each randomly sampled network in (A). Note that edges were defined for neighboring tissue-spots only, thus the maximum degree for a given spot is 6. We observed an increased average degree with HFD feeding, indicating more highly-localized LAM expression. **(C)** The distributions of (A) and (B) over time. **(D)** The mean clustering coefficient [47] in LAM networks compared with the optimal hard threshold (OHT) [21] of the graph's adjacency matrix **A**. The clustering coefficient is the network average of the fraction of pairs of a node's neighbors that are connected. Clustering coefficient is one for a fully connected graph but tends to zero on a random graph as the graph becomes large. The OHT is used to estimate the true rank of the graph's adjacency matrix **A**.

623 ## Supplementary Tables

| Time | n Cells | Mean Area ($\mu$m) | STD Area ($\mu$m) | Mean Diameter ($\mu$m) | STD Diameter ($\mu$m) |
|---|---|---|---|---|---|
| ND | 378 | 8,704 | 5,040 | 52.63 | 40.05 |
| HFD8 | 663 | 13,004 | 5,853 | 64.33 | 43.16 |
| HFD16 | 299 | 19,774 | 10,954 | 79.33 | 59.05 |

Supplementary Table 1: **Mean adipocyte size with HFD feeding.** Adipocyte area distributions measured in images from high-resolution microscopy.

| Cell Type | ND | 8w | 14w |
|---|---|---|---|
| B cell | 362 | 282 | 183 |
| Dendritic cell | 143 | 1,058 | 882 |
| Mac1 | 136 | 317 | 510 |
| Mac2 | 25 | 406 | 36 |
| Mac3 | 179 | 519 | 57 |
| Mac4 | 15 | 613 | 1,411 |
| Mac5 | 4 | 333 | 1,870 |
| Monocytes | 175 | 714 | 1,009 |
| NK cell | 96 | 505 | 125 |
| Other | 33 | 78 | 54 |
| T cell | 93 | 1,298 | 299 |

Supplementary Table 2: **Number of cells by type at each diet condition.**

| Cell Type | ND vs. 8w | ND vs. 14w | 8w vs. 14w |
|---|---|---|---|
| B cell | 0.000000 | 0.000000 | 0.000000 |
| Dendritic cell | 0.000000 | 0.000000 | 0.000000 |
| Mac1 | 0.000000 | 0.028024 | 0.000000 |
| Mac2 | 0.000000 | 0.000000 | 0.000000 |
| Mac3 | 0.000000 | 0.000000 | 0.000000 |
| Mac4 | 0.000000 | 0.000000 | 0.000000 |
| Mac5 | 0.000000 | 0.000000 | 0.000000 |
| Monocyte | 0.000000 | 0.680789 | 0.000000 |
| NK cell | 0.632164 | 0.000000 | 0.000000 |
| Other | 0.000000 | 0.000000 | 0.000000 |
| T cell | 0.000000 | 0.000000 | 0.000000 |

Supplementary Table 3: **T-tests of edge weight distributions between diet-conditions.** Results for Welch's t-tests between the edge distributions of networks constructed from the entire tissue-capture area. Edge weights are defined as the harmonic mean of predicted CARD proportions between neighboring tissue-capture spots. With $\alpha = 0.01$ (Bonferonis $\hat{\alpha} = 0.0003$).

| Time | Cell Type | Protein | $p$-value | Fold Change (log2) |
|---|---|---|---|---|
| ND | Macrophages | CD11b | $< 0.0001$ | 0.963 |
| ND | Macrophages | F4-80 | $< 0.0001$ | 0.996 |
| ND | Monocytes | CD11b | $< 0.0001$ | 1.200 |
| ND | B cells | CD19 | $< 0.0001$ | 1.977 |
| ND | NK cells | CD4 | $< 0.0001$ | 0.618 |
| ND | NK cells | CD3 | $< 0.0001$ | 0.928 |
| 8w | Macrophages | CD4 | $< 0.0001$ | 0.520 |
| 8w | Macrophages | CD11b | $< 0.0001$ | 1.452 |
| 8w | Macrophages | F4-80 | $< 0.0001$ | 1.252 |
| 8w | Macrophages | Mac-2 | $< 0.0001$ | 0.619 |
| 8w | T cells | CD3 | $< 0.0001$ | 0.588 |
| 8w | B cells | CD19 | $< 0.0001$ | 1.220 |
| 14w | Macrophages | CD11b | $< 0.0001$ | 1.410 |
| 14w | Macrophages | F4-80 | $< 0.0001$ | 1.441 |
| 14w | Macrophages | Mac-2 | $< 0.0001$ | 1.413 |
| 14w | T cells | CD3 | $< 0.0001$ | 0.526 |
| 14w | B cells | CD19 | $< 0.0001$ | 0.699 |

Supplementary Table 4: **Protein Validation of Predicted Cell Types.** Results of Wilcoxon rank-sum tests for differential protein expression for each cell type against all other cells at each time point. We adjusted $\alpha$ using Bonferroni's correction and required that the fold change (log2) was greater than 0.5. Using this conservative criteria we show that our cell type annotations are highly aligned with protein expression.

| Cell Type 1 | Cell Type 2 | Number Up | Number Down |
|---|---|---|---|
| Mac1 | Mac2 | 374 | 141 |
| Mac1 | Mac3 | 375 | 339 |
| Mac1 | Mac4 | 184 | 141 |
| Mac1 | Mac5 | 217 | 352 |
| Mac1 | Monocytes | 394 | 119 |
| Mac2 | Mac3 | 89 | 234 |
| Mac2 | Mac4 | 170 | 413 |
| Mac2 | Mac5 | 208 | 932 |
| Mac2 | Monocytes | 136 | 68 |
| Mac3 | Mac4 | 306 | 356 |
| Mac3 | Mac5 | 485 | 822 |
| Mac3 | Monocytes | 145 | 59 |
| Mac4 | Mac5 | 46 | 52 |
| Mac4 | Monocytes | 210 | 77 |
| Mac5 | Monocytes | 546 | 288 |

Supplementary Table 5: **Number of differentially expressed genes between macrophage subtypes.** Results of pairwise differential expression analysis on genes expressed in at least 10% of the macrophage/-monocyte cell population between macrophage subtypes and monocytes. DEGs were identified using a non-parametric Wilcoxon rank sum adjusted using Bonferroni correction ($\alpha = 0.01$). We count the number of genes with fold change (log2) greater than 1 (up) and less than 1 (down).
